# Beta-Hydroxybutyrate but not NMN supplementation mimics caloric restriction reducing early mortality in *Daphnia*

**DOI:** 10.1101/2025.05.21.655400

**Authors:** A. Catherine Pearson, Lev Y. Yampolsky

## Abstract

NAD+ homeostasis is an important determinant of lifespan and may be a key mechanism of caloric restriction (CR) expansion of lifespan. Ketone bodies such as beta-hydroxybutyrate (BHB) that regulate NAD+ abundance and NAD+ precursors such nicotinamide mononucleotide (NMN), as are known to extend life in experimental animals and ameliorate age-related conditions in humans. We tested the hypothesis that chronic BHB and NMN exposure separately or in combination can extend lifespan in a model organism *Daphnia,* a freshwater zooplankton crustacean with the magnitude similar to that of the CR treatment. We also measured fecundity, lipofuscin accumulation, and lipid investments into offspring in *Daphnia* fed the full diet, full diet with BHB, NMN, and combined treatments, and fed the CR diet (25% of the full diet). We also conducted an RNAseq experiment comparing the two diets and the two exposure treatments. We show that BHB exposure, but not NMN exposure reduces early life mortality in *Daphnia* fed the full diet to levels similar to those observed under CR without compromising fecundity. We also observed that in a combined exposure cohort, NMN nearly eliminates the beneficial effect of BHB. None of the treatments affected lipofuscin accumulation, but the NMN and the combined treatment mimicked the effect of CR on neonate size in older females. We show that BHB-treated *Daphnia* change expression of a variety of genes, including genes with known longevity extending effects, but differential expression of few genes is consistent with the effects of CR and their functionality is not clear.

## Introduction

Among several types of metabolic interventions aiming to extend lifespan and health- span those that act through mimicking caloric restriction (CR) have emerged as particularly promising (Madeo et al. 2014; Longo et al. 2015; Gonzalez-Freire et al. 2020). There are several pathways through which naturally occurring metabolites can mimic CR, but two stand out in particular: carbohydrate and fat metabolism regulating insulin signaling pathway and activation of NAD+-dependent sirtuins with convincing data on lifespan extension by activation of these pathways available in yeast, *Caenorhabditis elegans,* and *Drosophila,* (Baur et al. 2012; Longo et al. 2015; Gonzalez-Freire et al. 2020). In this report we explore life- extending effects of two key metabolites - beta-hydroxybutyrate (BHB, also often abbreviated as βOHB) and nicotinamide mononucleotide (NMN), a key precursor of NAD+, capable of regulating one or both pathways to the emerging longevity genomics model organism, the plankton crustacean *Daphnia.*

BHB is one of the key ketone bodies that naturally accumulate in animals maintained on ketogenic diet, which has been proven effective in promoting longevity (Balietti et al. 2010; Edwards et al. 2014; Campisi et al. 2019; Han et al. 2020; Wang et al. 2021) through serving as an energy source alternative to carbohydrates and by performing crucial signaling functions. As an energy source, ketone bodies differ from glucose in their energy metabolism propensity to increase the NAD+/NADH ratio because production of acetyl-CoA from BHB consumes half the amount of NAD+ molecules, mole to mole, consumed by production of acetyl-CoA by glycolysis and pyruvate oxidation (Newman & Verdin 2017). Besides being an alternative energy source, BHB can also play a variety of signaling roles (Newman and Verdin 2017; Puchalska & Crawford 2021; Wang et al. 2021; Nelson et al. 2023), including, notably, activation of NAD-dependent sirtuins by increasing NAD+ availability and inhibition of class I histone deacetylases (HDACs), thereby maintaining elevated gene expression. NAD- dependent sirtuins are mitochondrial protein deacetylases (Stein & Imai 2012) that function as nutrient-responsive regulators and are thought to extend lifespan in yeast, worms, and flies and to be one of the underpinnings of CR extension of lifespan (Sugishita et al. 2024).

Histone hyperacetylation induces expression of several key regulatory proteins, including Foxo3a, the mammalian ortholog of the stress-responsive transcriptional factor DAF-16 that regulates lifespan in *C. elegans* (Shimazu et al. 2013; Newman and Verdin 2017). Another downstream process affected by histone hyperacetylation is autophagy (Yi et al. 2012).

In parallel, studies focused on the role of NAD+ homeostasis in organisms like yeast (Anderson et al. 2003) and *Drosophila* (Balan et al. 2008) have identified NMN as the key precursor of NAD+ and as a promising molecule for promoting longevity and metabolic health (Madeo et al. 2014; Imai & Guarente 2016; Rajman et al. 2018; Yoshino et al. 2018). In mice, the NMN supplementation has been shown to enhance NAD+ levels in multiple tissues, improving mitochondrial function, insulin sensitivity, and overall metabolic health (de Picciotto et al. 2016; Mills et al. 2016). Increasing NAD+ availability through NMN supplementation has been linked to the activation of sirtuins (Imai & Guarente 2016), particularly SIRT1, a well-known regulator of cellular stress resistance and metabolic adaptation. SIRT1 activation mimics several effects of CR, including enhanced mitochondrial biogenesis, reduced oxidative stress, and improved DNA repair (Gomes et al. 2013; Sinclair & Guarente 2014; Cantó et al. 2015;). Given the shared role of both BHB and NMN in increasing NAD+ availability and promoting sirtuin activation, it is hypothesized that both compounds may act synergistically to mimic the beneficial effects of CR on lifespan and healthspan.

*Daphnia* (Crustacea: Branchiopoda) is a classic model organism for longevity studies, one of the first animal in which the lifespan extending effect of CR has been demonstrated (Ingle 1933). *Daphnia* are particularly suitable for the studies of environmental effects on longevity because its mode of reproduction, cyclic parthenogenesis, allows placing outbred genetically identical individuals into different environments, e.g., on different diets, eliminating confounding of environments with genotypes, within-cohort heterogeneity, or the need to use highly inbred lines. More recent studies extensively documented diet effects on longevity in *Daphnia* (Lynch and Ellis 1983; Lynch 1989; Pietrzak et al. 2010; Nguyen et al. 2021; see Lampert 2025 for a review), including evidence that the life-extending effect of CR is only present in long-lived, but not in short-lived genotypes (Beam et al. 2024). With this in mind, testing for the possible CR mimicking effect of BHB and NMN in *Daphnia* could be particularly instructive, as it might elucidate mechanisms of further extending healthy lifespan beyond the genetically determined benchmark.

In *Daphnia,* just as in many other organisms (Holliday 1989; Shanley & Kirkwood 2000; Mc Auley 2022), the longevity extension on calorically restricted diet is accompanied by drastically reduced reproduction rate, raising the questions about the role of reproduction vs. maintenance trade-off in longevity in general and in the mechanism of CR effect in particular. It would be particularly interesting to observe a treatment that would mimic the CR effect without sacrificing reproduction, thus escaping the apparent trade-off, something that is observed, for example, in nereiid flies under certain combinations of environmental conditions (Adler et al. 2013) or in sterol-supplemented high protein diet-fed *Drosophila* (Zanco et al. 2021; Piper et al. 2023). Furthermore, the effect of CR on reproduction in *Daphnia* is not limited to just production of fewer offspring. Counterintuitively, *Daphnia* are known to produce fewer, but larger and better provisioned offspring under dietarily restricted conditions (Guisande & Gliwicz 1992; Boersma 1995 Mckee & Ebert 1996). Any CR-mimicking intervention may as well mimic this resource allocating strategy, elucidating its possible mechanisms.

Besides fecundity and resource allocation to offspring, both CR and CR-mimicking metabolic interventions are likely to alter other age-related changes in *Daphnia* physiology, possibly revealing the mechanistic explanation for longevity extension. We have recently demonstrated that aging *Daphnia* accumulate lipofuscins, products of oxidative damage to lipids and proteins easily detectable in vivo due to *Daphnia*’s transparent tissues (Lowman and Yampolsky 2023). Thus one of the goals of this study is to evaluate the degree of lipofuscin accumulation in full diet control relative to CR and CR-mimicking treatments.

Finally, we aimed to utilize genomic resources available in *Daphnia* to expose the underlying transcriptional changes as the basis of both CR and CR-mimicking metabolic interventions by an RNAseq experiment. In doing so we pursued two goals: first, to functionally characterize transcriptional changes in response to CR, and to BHB and NMN exposure separately; second, to detect specific gene expression changes in response to metabolic interventions that are collinear with those in response to CR. The second goal would not provide a proof of the mechanism of CR mimicking, but will allow to exclude genes whose changes are coincidental with the transcriptional response to CR.

## Materials and Methods

### Clone’s provenance and maintenance

The experiment was conducted using two parthenogenetically maintained isolates of *D. magna,* GB-EL75-69 and IL-M1-8, previously characterized as a long-lived and a moderately long-lived genotypes, respectively (Beam et al. 2024). These clones have been originally extracted, respectively, from a permanent lake in London, UK, and an intermittent summer-dry pond in Jerusalem, Israel. The two clones will be referred to thereafter as the GB and IL clones. Both have been obtained from Basel University Daphnia stock collection (Basel, Switzerland) and maintained in the lab for over 5 years in modified ADaM zooplankton medium (Klüttgen et al. 1994; https://evolution.unibas.ch/ebert/lab/adam.htm) at 20 °C on the diet of the green alga *Scenedesmus acutus* (synonym: *Tetradesmus obliquus*) grown in the lab on constant light in Jüttner et al. (1983) algal medium. To obtain animals for the experimental cohorts, three juvenile females of each clone were taken out of the stock cultures and maintained individually in 20 mL of ADaM medium, fed with 2E5/mL *Scenedesmus* cells per day, thus initiating three replicate cohorts. Second or thirds clutch offspring of these progenitor grandmothers were collected within 24 hours of birth and maintained in 100 mL glass jars under the same conditions, serving as mothers of the experimental animals.

### Experimental design

Offspring of the maternal generation were collected from mothers in each of replicate and each of the two clones in the same manner and split, within 24 hours of birth, into *ad libitum* treatment receiving 4E5/mL *Scenedesmus* cells per day and CR treatment, fed 1E5/ml *Scenedesmus* cells per day. These two dietary treatments will be hereafter referred to as Control400 (full diet) and Control100 (CR diet). These individuals were kept in 100 mL glass jars until the age of 6-7 days, at which point they were transferred, in groups of 5, into 50 mL plastic tubes with the bottoms cut off and replaced by 1mm nylon. These tubes were placed into racks, 32 tubes per rack, and each rack was placed into a 4L glass aquarium filled with ADaM medium to achieve 20 mL volume per *Daphnia.* Food has been added to these aquaria daily with the concentration of 1E5 and 4E5 cells per mL in the two diet treatments and the medium was replaced every 4 days. Census was taken every other day and medium volume and food quantity was adjusted to account for attrition in the cohorts.

Racks containing tubes with *Daphnia* were removed from the holding aquaria for 2 hours every day and placed, after rinsing in fresh ADaM medium, into trays containing either 2.5 mM BHB, or 0.1mM NMN solutions in ADaM, or a solution of both chemicals together, at the same concentrations. Control cohorts in each of the diet treatments were placed for the same amount of time into identical trays containing ADaM medium. Thus, individuals from the two clones shared the exposure handling and solution, and all treatments shared the medium and food outside of exposure times. This resulted in the following treatments: Control400 (full diet), full diet with BHB exposure, full diet with NMN exposure, full diet with exposure to both BHB and NMN, and Control100 (CR diet).

Age-specific mortality data were analyzed for differences between high food controls and CR, BHB, and NMN treatments using Kruskal-Meyer survival analysis, proportional hazard model and parametric survival model assuming log-logistic distribution. The latter model is particularly suitable for the observed survival curves due to non-linear changes in log-hazard with age and high sensitivity to early and late-life mortality (Muse et al. 2021).

### Fecundity measurements

To measure fecundity females carrying a brood of embryos in late stages of development (single black eye stage, < 24 hours before brood release, whenever possible) were sampled from the experimental tanks and kept in individual vials containing 20 mL of the medium, with the amount of food added daily matching their treatments, until the brood was released and the neonates could be counted. Six females were sampled from each combination of clone and treatment. After brood release the females were returned to the inserts from which they were sampled. Such fecundity counts were conducted 6 times throughout the lifespan at ages (SD) 13 (2.4), 34 (2.2), 52 (2.4), 86 (2.8), 112 (2.5), and 131 (2.3) days. To account for skipped clutches (0 eggs produced during an intermolt period, a frequent occurrence at older ages), simultaneously with sampling the females from the inserts, all females in this insert were screened, and clutch size of those with empty brood chambers and no visible ovaries was recorded as 0.

### Lipids provisioning and body length in oAspring

At the ages of 52 and 86 days the neonates released in fecundity measurement vials all treatments except the double-exposure treatment, were sampled from the broods produced by 3-6 replicate females per clone per treatment, up to 4 per replicate (fewer for smaller broods), within 18 hours after the brood release. These neonates were stained with Nile Red dye (2 hour staining in 1 ug/mL solution in ADaM medium, 2-4 neonates per 1 mL. After the staining the neonates were washed in ADaM and photographed under fluorescence Nikon Labophot 2 microscope using B-2A filter cube (excitation 450-490 nm, 505 nm dichroic mirror and 520 barrier filter), with the ND2 filter applied, a 4x lens and a 5 MP Moticam color camera. The amount of yellow-green fluorescence specific to lipid droplets in the resulting 8- bit images was they measured using ImageJ by calculating portion of pixels above the cut-off value of 150 fluorescence units (Supplementary Fig. 1A). For graphic representation purposes these estimates were then averaged over all neonates from the same clutch. For the 3-way ANOVA with treatments, clones and age classes as factors the clutch ID was included as a random effect nested within clones. None of age class interactions were significant and thus were removed from the model without affecting significance level of other main effects. The same neonates were also photographed in bright field and body length from the top of the head to the base of the caudal spine was measured in ImageJ.

**Fig. 1.**
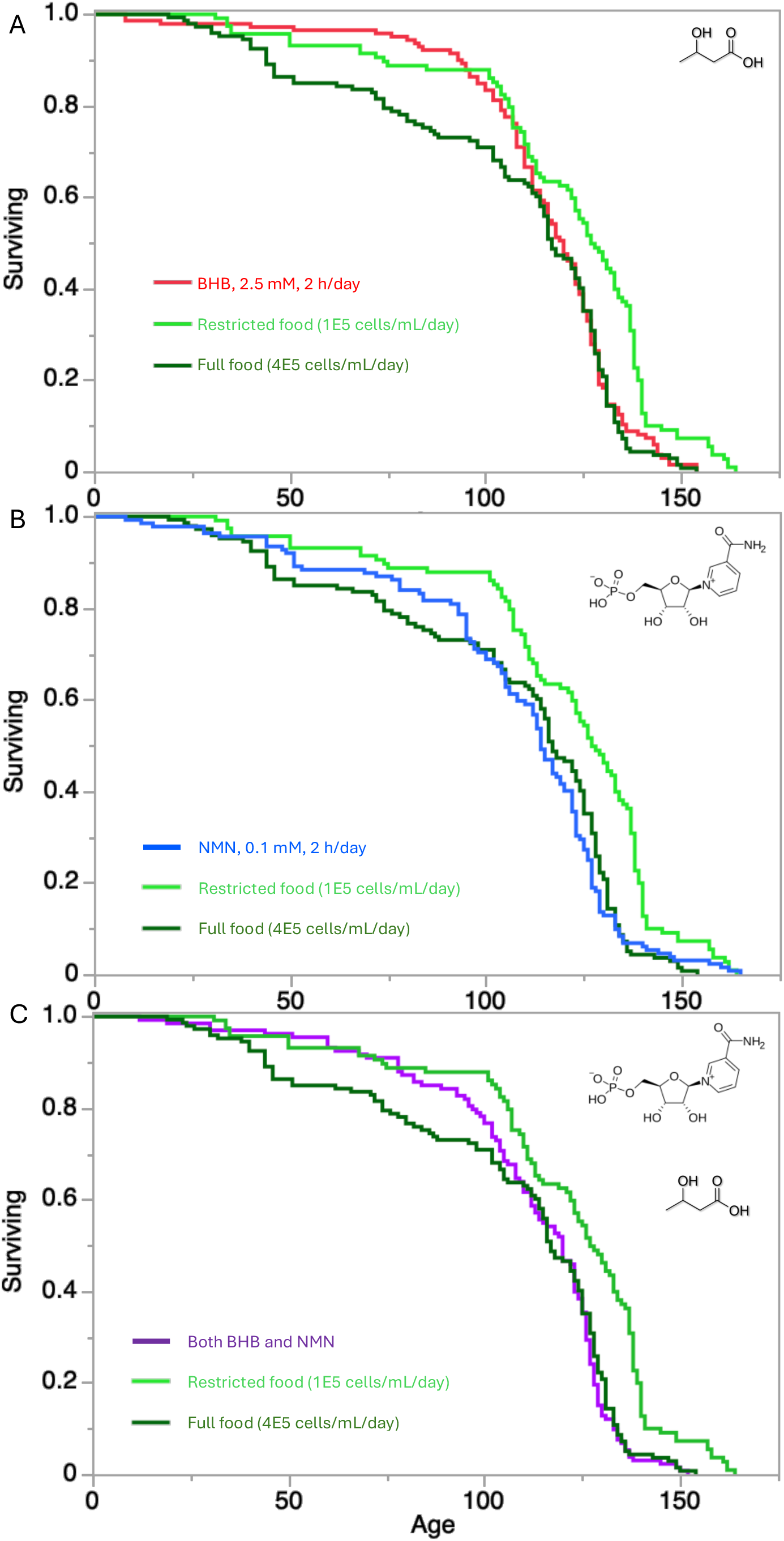
Survival curves of cohorts of *D.magna* on full diet (4E5 cells/mL/day; dark green), on restricted diet (1E5 cells/mL/day; light green) and treated with either BHB (A, red curve), or NMN (B, blue curve), or both (C, purple curve). See Table 1 for parametric survival analysis. Data from the two clones combined. See Supplementary Fig. 1 for survival curves for each clone separately.

**Table 1.** Parametric survival analysis with long-logistic model fit (log-likelihood ratio test).

| Source | DF | L-R $\chi^2$ | P |
| --- | --- | --- | --- |
| Contrast: All treatments |  |  |  |
| Treatment | 4 | 17.4 | 0.0016 |
| Clone | 1 | 37.16 | <0.0001 |
| Clone*Treatment | 4 | 7.09 | 0.13 |
| Contrast: caloric restricted diet vs. full diet control |  |  |  |
| Treatment | 1 | 10.15 | 0.0014 |
| Clone | 1 | 9.18 | 0.0024 |
| Clone*Treatment | 1 | 3.66 | 0.06 |
| Contrast: 2.5 mM BHB vs. full diet control |  |  |  |
| Treatment | 1 | 5.1 | 0.024 |
| Clone | 1 | 18.33 | <0.0001 |
| Clone*Treatment | 1 | 2.66 | 0.1 |
| Contrast: 0.1 mM NMN vs. full diet control |  |  |  |
| Treatment | 1 | 0.03 | 0.87 |
| Clone | 1 | 23.35 | <0.0001 |
| Clone*Treatment | 1 | 0 | 0.99 |
| Contrast: BHB and NMN treatment vs. full diet control |  |  |  |
| Treatment | 1 | 1.24 | 0.27 |
| Clone | 1 | 20.37 | <0.0001 |
| Clone*Treatment | 1 | 0.95 | 0.33 |

### Lipofuscins accumulation and background autofluorescence and

To assess lipofuscins accumulation in aging *Daphnia* (Lowman and Yampolsky 2023) 6 replicate females per clone per treatment were sampled at the age of 115 (3.1) days and autofluorescence was measured in the same manner as above. Three images were taken from each female: nephridium, mid-body to exclude nephridium include thoracal fat body and abdomen (Supplementary Fig. 1B-D), with the abdomen image analyzed for to ROIs: abdominal fat boy and ovary. The images were analyzed in ImageJ by saving the histogram of the green channel (Supplementary Fig. 2) and calculating the portion of each ROI with fluorescence 25 units or more above the background mode. This background-subtracted cut- off corresponded to lipofuscin lumps emitting higher than the background (Supplementary Fig. 1C,D). Additionally, visual examination of fluorescence intensity histograms has lead to the observation that treatments differed in abundance of pixels with green fluorescence above 100 fluorescence units, without subtracting the background fluorescence (i.e., elevation of the right-hand shoulder of a typical distribution of background intensity). This value was used as an arbitrary cut-off to evaluate differences among treatments in dispersed (background) fluorescence. This quantity may be interpreted as a measure of abundance of oxidized forms flavins in tissues. Flavins, along with NAD(P)H are the main endogenous fluorophores and whose fluorescence may serve as an indicator of the intracellular redox state (Schneckenburger & König 1992; Monici 2005). NADH fluorescence is unlikely to contribute to the measured background autofluorescence given the excitation wavelength used (chosen to quantify lipofuscins autofluorescence rather than the background intensity and to reduce UV-induced oxidative stress that can skew the measurements, Monici 2005).

**Fig. 2.**
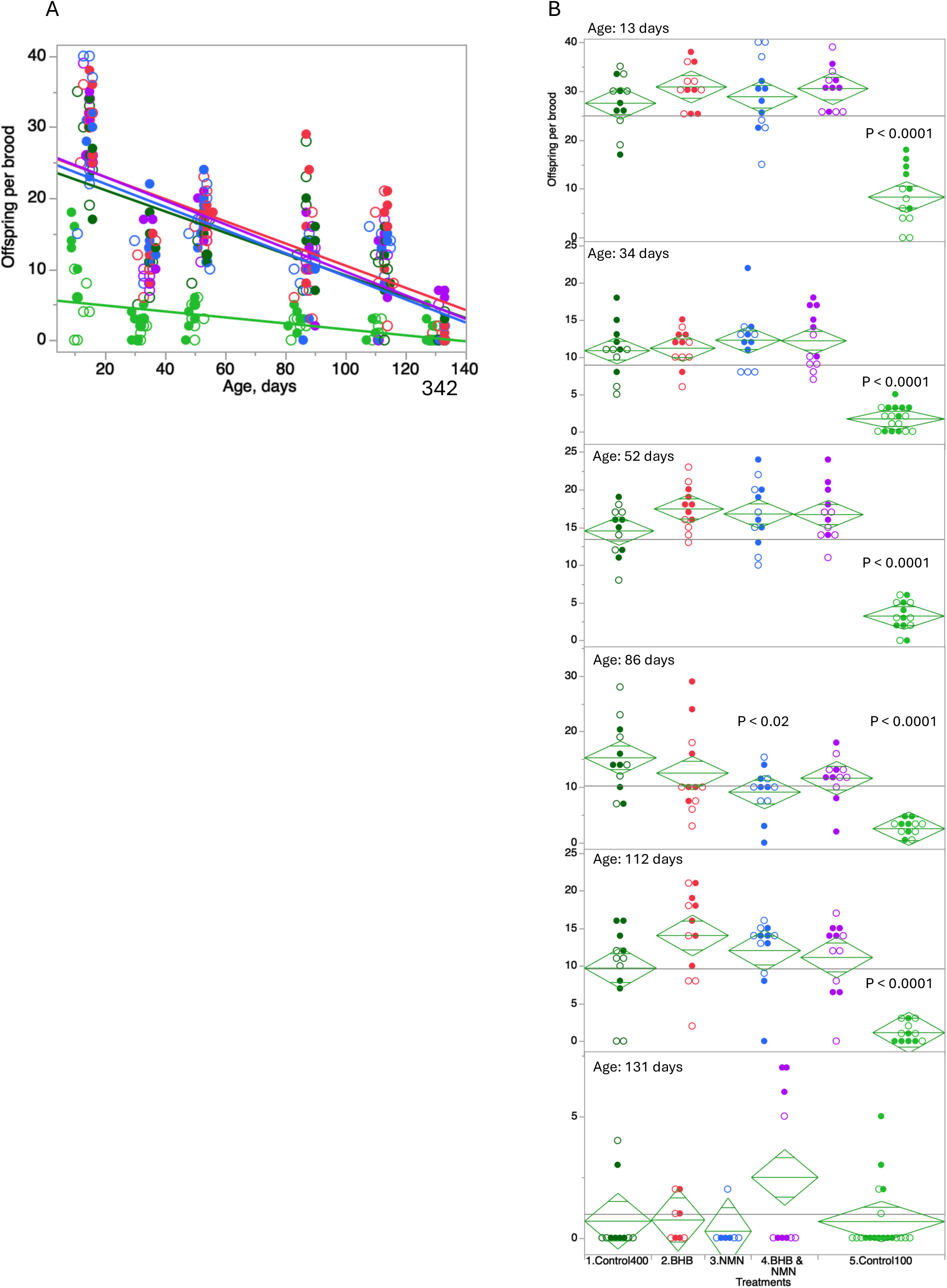
Fecundity (oNspring per brood) in 2 clones of *D. magna* in 6 age classes under each of the 5 treatments. Closed circles: GB clone, open circles: IL clone. Dark green: full diet control, light green: CR control, red: 2.5 mM BHB, blue: 0.1 mM NMN, purple both treatments. The height of diamonds represents 95% CI around means; horizontal lines within the diamonds show significant overlap between groups (equivalent to pairwise t-test); the width of the diamonds is proportional to sample size. P-values: Dunnett test against the full diet control, shown where at least one comparison is significant. A: All data, linear regression lines drawn through all points within treatment. B: Each age class separately.

**Fig. 3.**
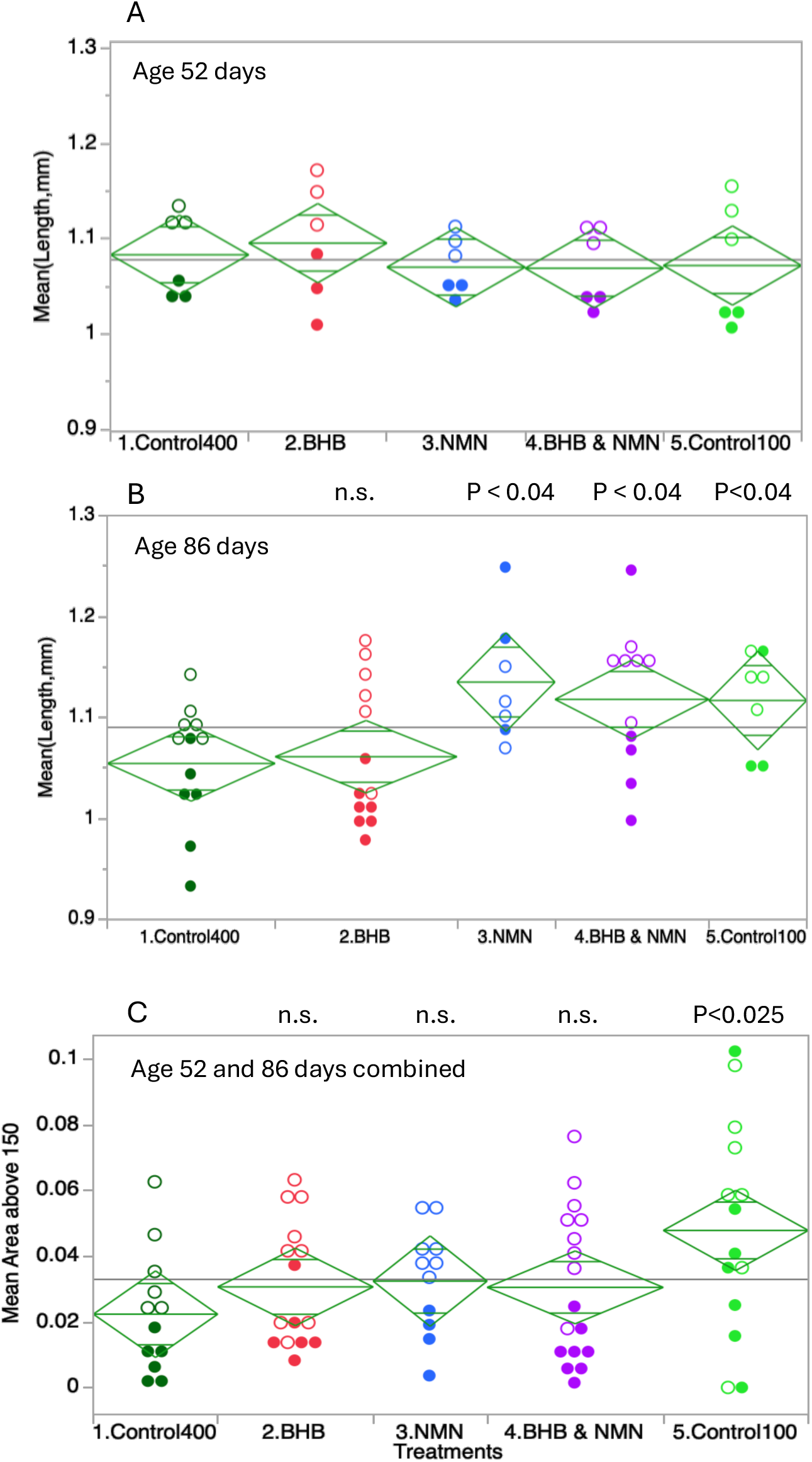
Neonate body length (A, B) and lipid content (portion of neonate body image occupied by lipid bubbles, Nile Red fluorescence intensity above 150 units, C); age classes combined because no significant diNerence between them (C). Each point is an average of several neonates from the same clutch. Symbols, colors and P-values as on Fig. 2. See Supplementary Table 3 for 2-way ANOVAs.

### RNAseq experiment

At the age of 75±2 days (i.e. at about 10% cohort mortality in the CR treatment and about 20% mortality in the full diet control) 3 females per clone were sampled from each of the control, CR, and BHB treatments and 2 females per clone from the NMN treatment, resulting in the total of 22 samples. Females were sampled at the same phase of the ovary/molting cycle by being selected within 24 hours after egg-laying. These freshly laid eggs were removed from the brood chambers and whole body somatic tissues of the females were homogenized in liquid nitrogen and total RNAs were immediately extracted using Qiagen RNeasy Plus Mini kit. RNA library preparation was performed using NEBNext Ultra II Directional RNA Library prep Kit (NEB, Lynn, MA) following the manufacturer’s protocols. The libraries were sequenced with Illumina Novoseq 6000, S4 flow cell, PE100. The quality check of Indexed sequences was performed by Fastqc, and indexed sequences were trimmed using an adaptor sequence by TrimGalore-0.4.5.

Sense reads were mapped to *D. magna* Xinb3 reference transcriptome (BioProject ID: PRJNA624896; D. Ebert and P. Fields, personal communication; Fields et al., in preparation) using STAR (Dobin et al. 2013) and genes with differential expression (DE) among treatments and between clones were identified using DEseq2 (Love et al. 2014). In DEseq2 analysis the replicates were used as a random variable nested within clones, except for the NMN vs.

Control contrast, where nested model could not be used due to only 2 replicates available for the NMN treatment; in this contrast the random variable was omitted from the model. DEseq2 P-values were adjusted for multiple tests by either built-in FDR procedure, for standalone Treatment vs. Control analysis, or, separately within only the set of genes with a significant DE between the two diet controls, in order to single out genes that responded to either BHB or NMN treatments matching a significant DE between CR and full diets. Genes with DE in response to CR, BHB and NMN treatments were functionally characterized using previously obtained GO annotation of *D.magna* transcriptome (Dua et al. 2024) using PANTHER 18.0 classification system (Mi et al. 2019).

To visualize transcriptional differences consistent with the effect of CR principal component analysis was conducted on TPM (transcripts per million reads) data on genes in which at there was a DE with |log2(FC)|>1 and padj<0.1. To pinpoint genes with expression pattern consistent with mortality differences between treatments the following additional criteria were used: 1) a significant difference between each BHB or NMN treatment and the full diet control as specified above and 2) mean TMP intermediate between the full diet and CR controls. Additionally, we specifically identified genes in which mean TPM was intermediate between the two controls in the IL clone, in which mortality difference between BHB and full diet control was the stronger of the two clones.

## Results

### Longevity and fecundity under caloric restriction and BHB and NMN supplementation

Exposure to 2.5 mM BHB for 2 hours per day reduced early mortality of females fed the full diet in both clones of *D.magna* to lower levels similar to those observed in the CR treatment (Fig.1 A). This effect was more pronounced in the shorter lifespan clone IL than in longer lifespan clone GB (Supplementary Fig S1). At around median lifespan full compensatory mortality was observed in the BHB treatment, reverting the survival curve back to that of the full diet control (Fig. 1 A). The proportional hazard test, being sensitive to late-life mortality failed to reach a significance level in a two-way, entire lifespan comparison of BHB-treated cohort to the full diet control cohort (log-likelihood c2= 2.77; P<0.1) and was only modestly significant for the IL clone when each clone was analyzed separately (log- likelihood c2= 4.09; P<0.05). However, the same comparison was highly significant when only the cohort’s age-specific mortality at ages below median lifespan were analyzed (log- likelihood c2= 10.17; P<0.0014). Thus, the BHB treatment increased longevity by reducing early-life mortality to mimic CR, with the magnitude of the effect different in the two genotypes tested.

The same effect of the NMN treatment was smaller in magnitude (Fig.1 B), only visible in the GB, but not IL clone (Supplementary Fig. S1) and not significant in the parametric survival test (Table 1). Furthermore, the NMN exposure seemed to reduce the BHB effect in the combined treatment with NMN and BHB (Fig.1 C; Supplementary Fig. S1).

Fecundity (offspring per brood) significantly decreased with age (Fig. 2, Table 2) and was significantly lower in the CR diet. A significant Treatment-by-Age interaction reflects the fact that loss of fecundity with age was significantly greater in the full diet that in the CR diet treatment. In most age classes the BHB, NMN, and combined treatments resulted in a slightly higher fecundity values than in the full diet control, none of which were significantly different from the full diet control in Dunnett test; at the age of 86 days (i.e. close to the age at which late-life reproductive resurrection often starts to be observed, Dua et al., 2024), in all three treatments fecundity was slightly lower than in the full diet control, with the NMN treatment reaching P<0.02, significant within that age class, but not significant after multiple test correction across all age classes. At the oldest age class there were some larger clutches observed in the CR treatment and in the combined BHB and NMN treatment, but overall there was no significant difference among treatments. Thus, BHB, NMN, and their combination did not reduce fecundity of full diet-fed *Daphnia* at any age.

**Table 2.** ANOVA of the effects of age (continuous variable), treatments, clones (categorical variables), and their interactions on *Daphnia* fecundity. . See Fig.2.

| Source | DF | Sum of<br>Squares | F Ratio | Prob > F |
| --- | --- | --- | --- | --- |
| Age | 1 | 11273.3 | 304.4 | <0.0001 |
| Treatments | 4 | 9035.0 | 61.0 | <0.0001 |
| Clone | 1 | 163.1 | 4.4 | 0.04 |
| Treatments*Clone | 4 | 75.7 | 0.51 | 0.73 |
| Treatments*Age | 4 | 1668.3 | 11.3 | <0.0001 |
| Clone*Age | 1 | 15.7 | 0.42 | 0.52 |
| Treatments*Clone*Age | 4 | 92.3 | 0.62 | 0.65 |
| Error | 343 | 12702.5 |  |  |

### Neonate length and lipid content

Females in the oldest age class (86 days) produced significantly larger neonates in low food than in low food, but this effect was not observed in younger females (52 days; Fig. 2 A, B; Table 2). In the older females this effect was fully mimicked by the BHB treatments to which high food females were exposed (Fig. 2B). No such mimicking was observed for the NMN treatment. There was also a significant between-clone effect with the IL females consistently producing larger offspring than the GB females. Similarly, in both age classes females produced neonates with higher lipid content in the low food treatment than in the high food treatment (Fig. 2C, Table 2), with the IL females also producing better provisioned offspring. Yet, there was no evidence that either NMN or BHB treatments mimicked that effect of CR on lipids provisioning to the offspring.

### Background and lipofuscins autofluorescence

We observed significant differences between clones and among treatments in right-side skew in background green autofluorescence (portion of image above 100 units) in abdominal fat body (Fig. 4 A; Supplementary Fig. S3 A; Supplementary Table S1). The high and low food controls were significantly different in a post-hoc Dunnett’s test, with the BHB and NMN treatments showing intermediate values between the high and low food controls, in an additive manner (Fig. 4 A). Unexpectedly, there was a significant food-by-body region interaction in background autofluorescence with the data from thoracal fat body showing the opposite direction of change (Fig. 4 A, C; Supplementary Table S2). A moderately significant effect of treatments was observed for background fluorescence in the nephridium (Fig. 4E, Supplementary Table S1) with none of the treatments reaching a significant difference from the high food control, and no significant differences were found for the background fluorescence in the ovary (Fig. 4G, Supplementary Table S1).

Abdominal fat body and the ovary showed a significant differences between the two clones tested in green background autofluorescence with the GB clone consistently showing higher right-hand shoulder of intensity distribution than the IL clone.

Subtracting the background mode eliminated any differences among treatments and differences between clones in all tissues except ovaries (Fig. 1 B, D, F, H; Supplementary Table S1). The same was observed when higher cut-off values were used (i.e. portion of ROI above 50 units of background-subtracted fluorescence; data not reported). Thus, there were no differences among treatments with respects to abundance of lipofuscin lumps.

### DiAerential gene expression

Functional characterization of genes with significant DE in response to CR Counts and lists of genes showing a significant up- or down-regulation in response to CR, BHB, and NMN treatments are shown in Supplementary tables S3 and S4. Due to a large number of GOs and pathways identified among the PANTHER annotation of genes responding to CR, and, thus, low number of such genes per GO or pathway category, none of the detected GOs were significantly enriched relative to the numbers of genes annotated with such terms in the experiment-wide reference (Supplementary Tabel S5). Among top GO categories showing up-regulation by CR there were proteasomal ubiquitin-dependent protein degradation, oxidoreductase activity, carboxylic acid transport, phospholipid and nitrogen-containing compounds catabolism, and regulation of MAPK pathway. Those with down-regulated transcription included extracellular structural proteins and (possibly digestive) metallopeptidases, among others.

Concordance between DE in response to CR and metabolic interventions 216 genes showed differential expression in response to the CR treatment with 141 of them also passing the |log2(FC)|>1 criterion (Supplementary Table S3). Much fewer genes significantly responded to BHB ad NMN exposure, 15 (14) and 4 (4), respectively (numbers in parentheses: after application of the |log2(FC)|>1 criterion). The 15 genes with a significant response to BHB were not random with respect to those responding to CR, with the overlap between the 2 lists of 9 genes (expected: 0.39; Fisher’s Exact Test P<0.0001; Supplementary Table S3). This result also stands after the application of the |log2(FC)|>1 criterion. None of the 4 genes responding to NMN exposure also responded to the CR treatment. These results are consistent with the PCA analysis (Fig. 5 A) and expression heatmap (Supplementary Fig. S6), showing a significant separation between full diet and CR treatments (along the PC-1) and between the two clones (along the PC-2), with a slight tendency, within each clone, of overall expression profile to be shifted in the direction of PC-1 effect of CR (Fig. 5 A).

**Fig. 4.**
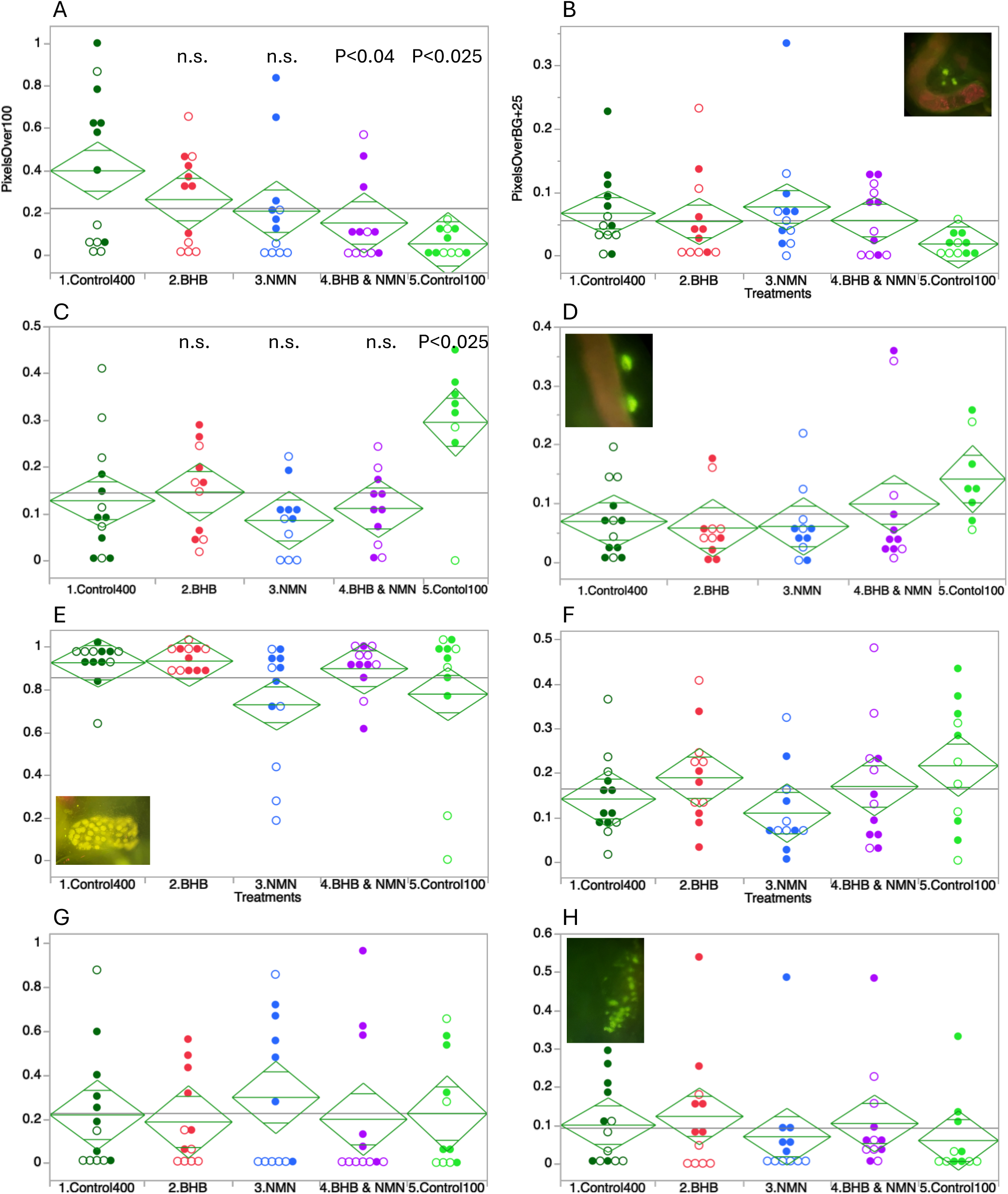
Late-age autofluorescence results. A, C, E, G: portion of green channel fluorescence above 100 units; B, D, F, H: portion of fluorescence 25 or more units above background mode. A, B: abdomen fat body, C, D: thoracal fat body; E, F: nephridium capsule; G, H: ovary. Closed circles: GB clone, open circles: IL clone. Diamond plot conventions and Dunnett test P-values as on Fig.2. See Supplementary Table S1 for 2-way ANOVAs, S2 for the analysis of Treatment-by-ROI interaction. Inserts: fragments of images of each ROI.

**Fig. 5.**
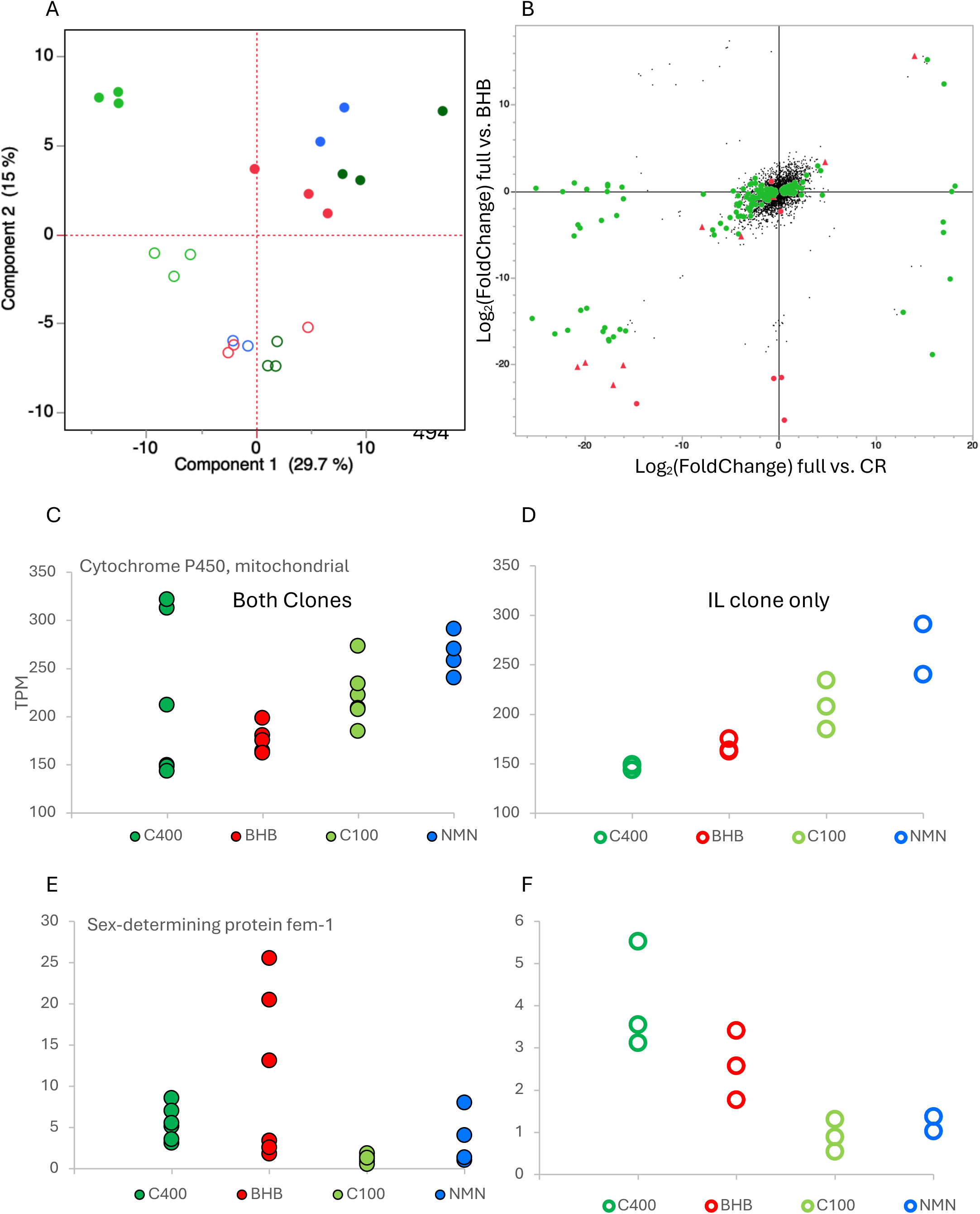
Differential expression of genes with a significant DE between full diet and CR control and between full diet control and BHB treatment. A: Principal component analysis on TPA values in genes with a significant DE between the two diets. B: “2D-volcano” plot showing log2(FoldChange) values between BHB treatments vs. the full diet control vs. log2(FoldChange) values between the two diets. Green circles: padi < 0.1 in the between-diets contrast; red circles: padi < 0.1 in BHB vs. full diet control contrast; red tringles: padi < 0.1 in both contrasts. Transcripts with no significant DE shown as black dots. See Supplementary Fig. S6 for analogous data for the NMN treatment. C – F: two genes meeting the criterion of significant DE in both contrasts and intermediate level of expression in the BHB treatment between the two diet controls. NMN data shown for a comparison. C, E: all data; D, F: data for the IL clone only. Colors and opens vs. closed circles on A and C – F as on previous figures.

Only two genes satisfied the criterion of the BHB treatment expression to mimic that in the CR treatment, a mitochondrial cytochrome P450 and a *Daphnia* ortholog of sex- determining protein fem-1 (Fig. 5 C – F). In both of them the effect was more pronounced in the IL clone.

## Discussion

We expected that, while in calorically restricted *Daphnia* BHB supplementation increases fecundity, but not lifespan, in *Daphnia* on an *ad libitum* diet BHB would extend lifespan, mimicking CR, without compromising fecundity. This is exactly what we observed, but only in the at earlier ages (Fig. 1A). Later in life, around median lifespan, mortality in BHB-treated *Daphnia* increased and the survival curve started to resemble that of the full diet control (Fig. 1A). We also expected a similar effect of NMN treatment. However, the life- extending effect of NMN was not significant (Fig. 1B) and in a combined NMN/BHB treatment nearly eliminated the early life mortality reducing effect of BHB (Fig. 1C).

The early mortality-reducing effect of BHB in a manner similar to that of CR is consistent with the role of BHB as a signaling molecule facilitating protective effects of NAD+ availability (Newman & Verdin 2017). However, lack of similar effect of NMN supplementation is inconsistent with this. Since the standard protocol for *Daphnia* lab maintenance implemented in this study (Goulden et al. 1982) implies the diet of algae grown with vitamin supplementation, nicotinamide included into the supplement may provide excess of NAD precursors, thus eliminating the effect of additional NMN supplementation.

An experiment with NMN supplementation on the background of B3 vitamin complex shortage would be needed to test this hypothesis.

Neither treatment provided an additional increase in fecundity relative to full diet (Fig. 2) with some evidence of reduced late-life fecundity in *Daphnia* supplemented with NMN (Fig. 2B). On the other hand, NMN treatment had an effect on late-life offspring size (Fig. 3B), mimicked the well-known effect of larger neonate size under limited diet (Boersma 1995, 1997; Mckee & Ebert 1996). Both treatments showed a trend towards the same with respect to neonate lipids provisioning, but no individual comparison to the full diet control was significant (Fig. 3 C).

**Table 3.** REML ANOVA of body size and lipid content of neonates, with Treatments, Clones, and maternal Age class as main effects (Age class interactions omitted where not significant). See Fig. 2 D. P < 0.01 shown in bold.

| Source | DF | DFDen | F Ratio | Prob > F |
| --- | --- | --- | --- | --- |
| Response: Neonate body length |  |  |  |  |
| Treatments | 4 | 57.3 | 1.567 | 0.20 |
| Clone | 1 | 57.4 | 43.817 | **<0.0001** |
| Treatments*Clone | 4 | 57.3 | 3.526 | 0.0125 |
| Age class | 1 | 57.4 | 4.136 | 0.05 |
| Treatments* Age class | 4 | 57.3 | 4.502 | **0.0031** |
| Clone*Month | 1 | 57.4 | 2.887 | 0.095 |
| Treatments*Clone* Age class | 4 | 57.3 | 1.495 | 0.22 |
| Response: Lipid content (portion of image covered by lipid bubbles) |  |  |  |  |
| Treatments | 4 | 53.2 | 4.243 | **0.0047** |
| Clone | 1 | 53.2 | 38.220 | **<0.0001** |
| Treatment*Clone | 4 | 53.2 | 0.724 | 0.58 |
| Age class | 1 | 53 | 1.899 | 0.17 |

The autofluorescence assays conducted on late-age females in this study had two purposes. One was to investigate the accumulation of lipofuscins known to be aging markers in a variety of organisms (Gray & Woulfe 2005.), including, with limitations, *Daphnia* (Lowman and Yampolsky 2023). The other goal was to assess red-ox state of tissues using oxidized flavins detectable by green autofluorescence as a proxy of red-ox potential shifted towards oxidation (Schneckenburger & König 1992; Monici 2005). In none of the four tissues sampled there was any difference in accumulation of lipofuscins (Fig. 4 B,D,F,H) either between full and restricted diet, or between any supplementation treatments and either of the controls. On the other hand, background fluorescence showed an unexpected opposite direction differences between the two diets in upper body and abdominal fat body, with the BHB and NMN treatments resulting in intermediate values in the abdominal fat body. We do not have an explanation for this observation other than perhaps abdominal fat body becoming less metabolically active in reproductively senescent females under restricted diet, thus resulting in less oxidative stress, which thoracal fat body continues to be metabolically active due to proximity to filtering and swimming appendages. *Daphnia* fat bodies are known to be the location of short-term fat reserves and of hemoglobin synthesis (Lampert 2025). If this is the case, the intermediate values of BHB and NMN treatments may also be interpreted as mimicking caloric restriction with respect to metabolism intensity.

A tendency to mimic caloric restriction is also observed in the overall transcriptional profiles of both clones studied for both BHB and NMN treatments. For the BHB treatment, in particular, there was a significant enrichment of genes that responded to both CR and BHB (9 such genes with expected <1). While higher than expected, the overlap was still small enough to draw any functional genomics conclusions. Genes showing down-regulation in both CR and BHB treatment included ankyrin repeats containing transposase, chitin deacetylase, as well as a number of uncharacterized proteins. A ribosomal protein gene and a gene encoding zinc metalloproteinase showed up-regulation in both. Only two genes with a significant differential expression between the two diets demonstrated intermediate transcriptional level in the BHB treatment (regardless of the |log2(FC)| for the BHB treatment), representing candidate genes for the BHB effects. These genes were those encoding a mitochondrial cytochrome P450 upregulated in both, and one of several *Daphnia* homologs of sex-determining protein *fem-1* known from nematodes and insects, which was downregulated in both CR and BHB treatment. Both genes met the criterion of intermediate expression in BHB only in one of the two clones sampled namely in the IL-M1-8 clone, which also showed a stronger reduction of early life mortality in the BHB treatment, consistent with the candidate status of these genes. Intriguingly, the two genes are linked, located in the same scaffold 146 kbp apart in the opposite orientation. In single cell transcriptomics data (Krishnan et al. 2024) the cytochrome P450 genes shows expression predominantly in one of the epithelial cell types, co-expressed with several cuticular proteins. It shares 33% sequence identity with its *Drosophila melanogaster* homolog, which is also known to be expressed in the epithelium, as well as in excretory tissues, larval gut, and, on the single cell level, in neurons. The function of the *Drosophila* homolog, similarly to other cytochromes, are reported to include oxidoreductase (monooxygenase) activity, possibly involving detoxification and antibacterial defense response (Ayres et al. 2009). The *fem-1* homolog (not detected in single cell data) has only a short area of homology with the *Drosophila* counterpart, which is characterized as an ankyrin repeat containing protein with predicted ubiquitin ligase activity expressed in male reproductive system and involved in protein metabolism regulation. It is not known whether the *Daphnia* protein in question has similar function, given that the area of homology is limited to the single ankyrin repeat and ankyrin repeat-containing proteins have a great variety of functionalities.

## Conclusions

BHB supplementation partially mimics caloric restriction by reducing early life mortality. Both BHB and NMN treatments mimic caloric restriction by reducing green autofluorescence indicative of oxidized flavins. Several genes showed coherent differential expression in CR and BHB treatments, but little functional conclusions could be made on the transcriptional basis of the observed effects.

## Acknowledgements

We are grateful to Patrick Bradshaw and Leonid Peshkin for useful discussions, to Ashit Dutta, Thomas Beam, Meridith Smith, Caroline Freeland, and Shelby Cutrell for laboratory assistance, and to Michael Shtutman and University of South Carolina Genomics center for RNAseq library preparation and sequencing. This work has been supported by Impetus Foundation grant to LYY.

## Data Availability

Raw reads are available at NCBI BioProject PRJNA1237747; Accessions SAMN47436000 - SAMN47436021; processed data in the form of read counts per transcript per library are available at NCBI GEO, Accession GSE292526. Life history data are available in the Supplementary Data file.

## Notes

### Competing Interest Statement

The authors have declared no competing interest.

