## Supplementary Tables and Figures for "Beta-Hydroxybutyrate but not NMN supplementation mimics caloric restriction reducing early mortality in *Daphnia*"

Clone: GB	Clone: IL

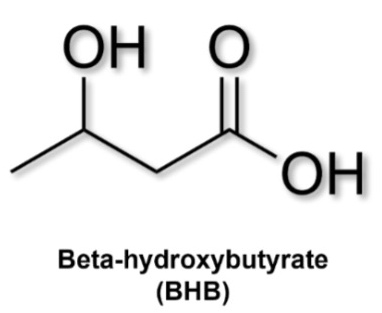

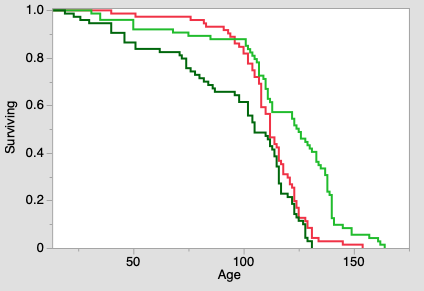

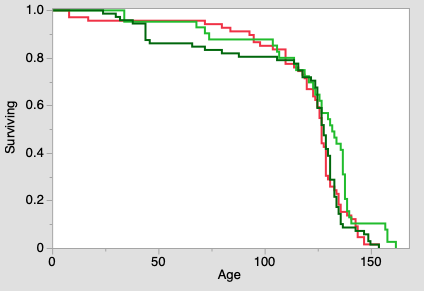

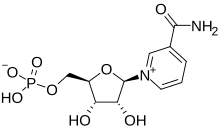

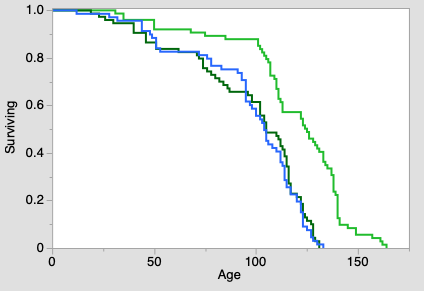

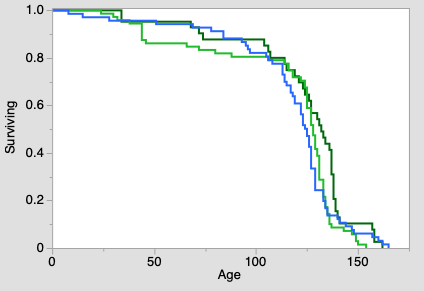

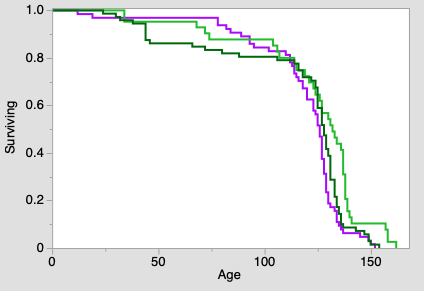

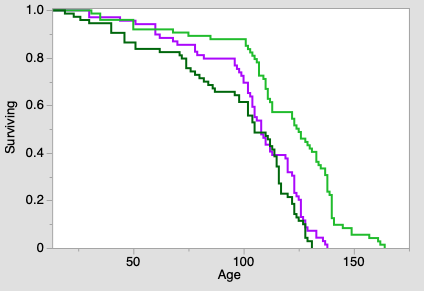

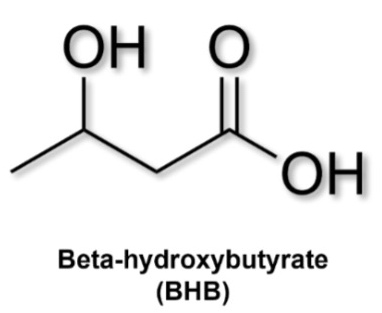

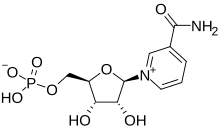

Supplementary Fig. S1. Survival curves of 2 clones of *D.magna* under conditions of full and restricted food (dark and light green) or full food with exposure to either BHB, or NMN, or both (red, blue, and purple curves).

A	B

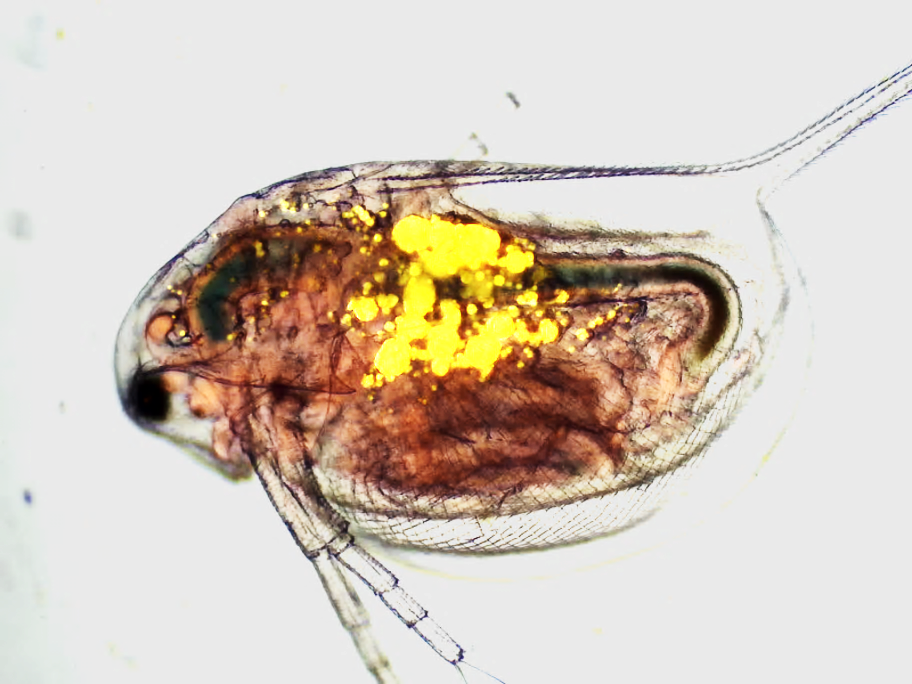

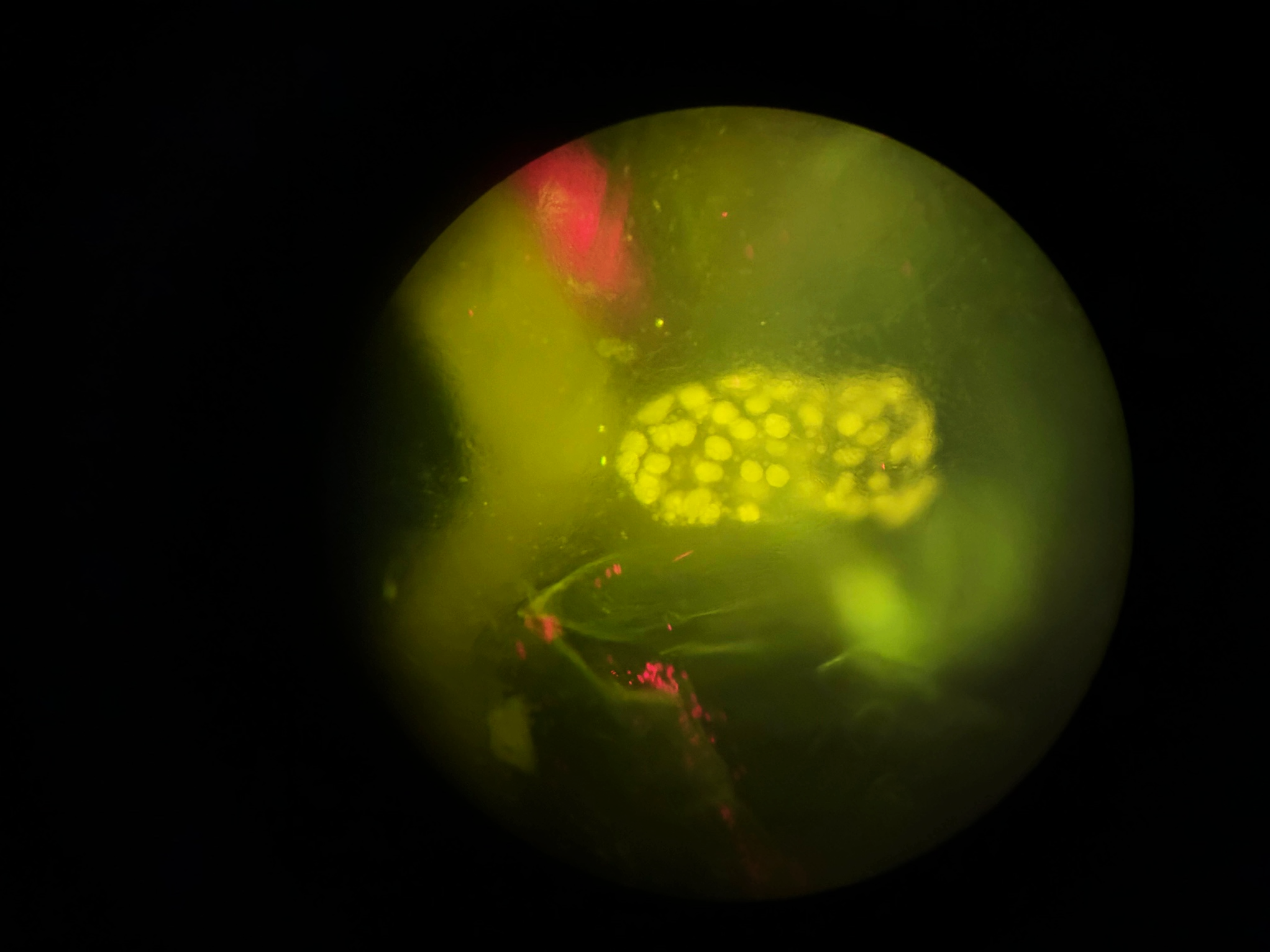

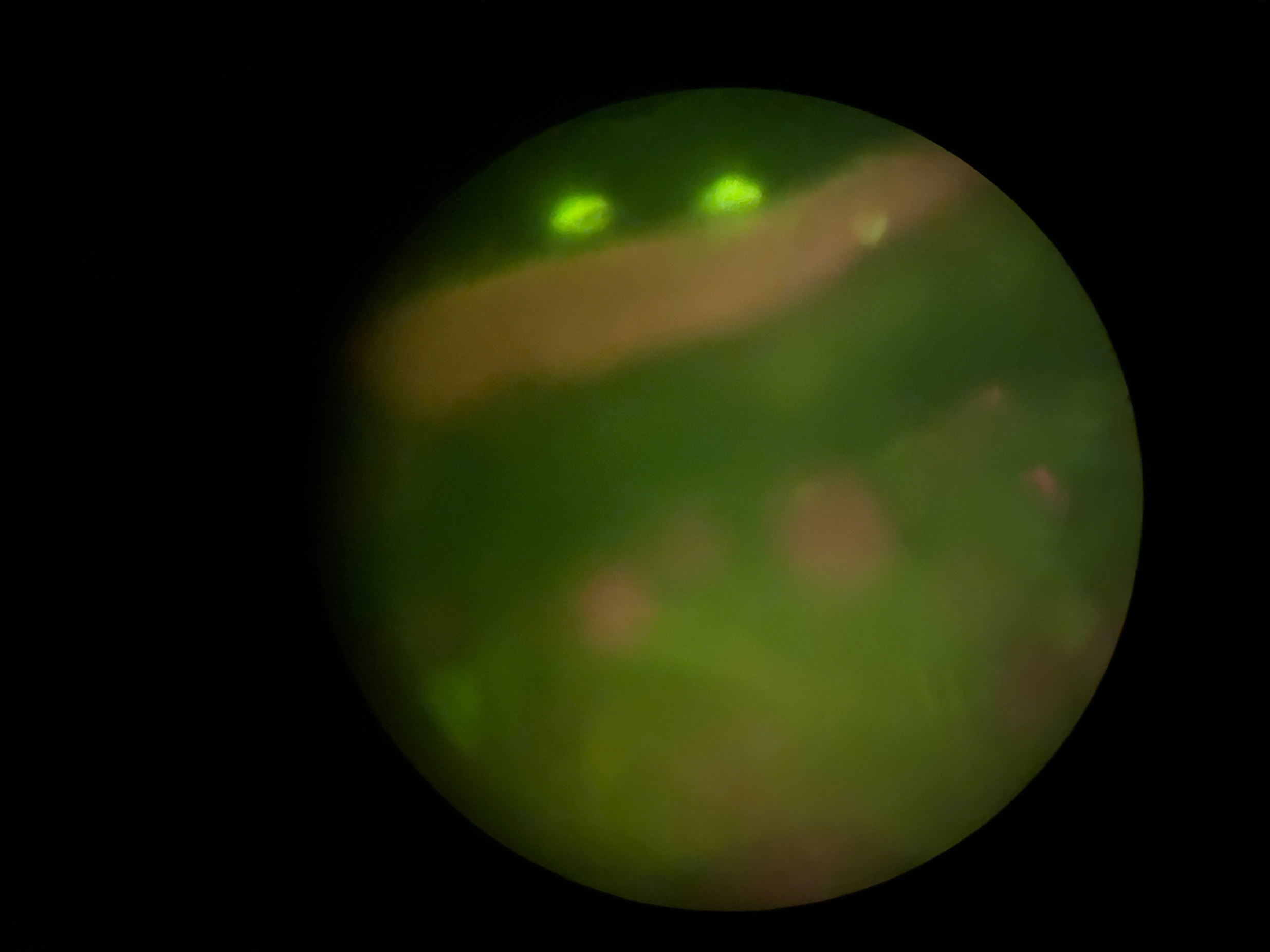

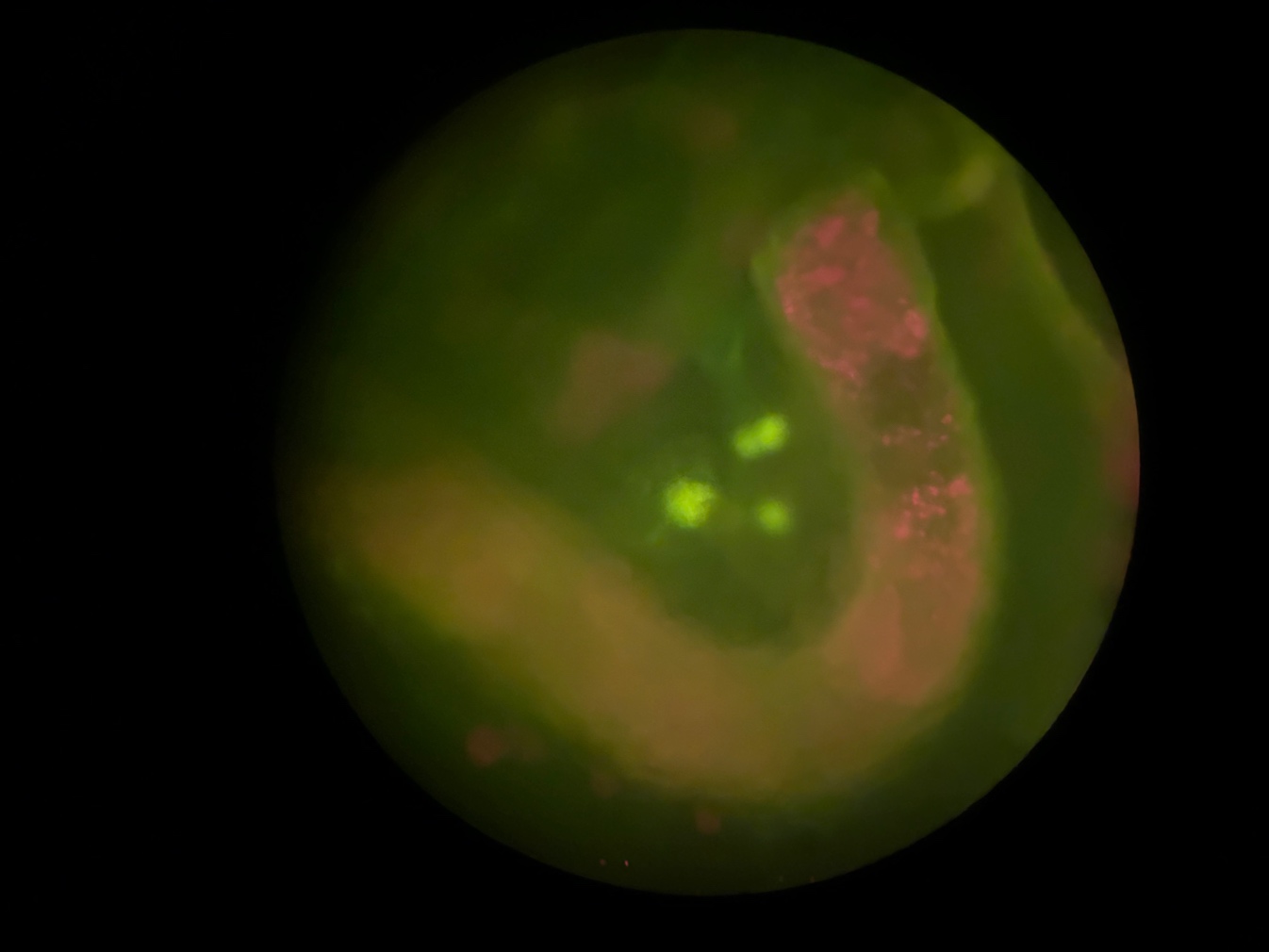

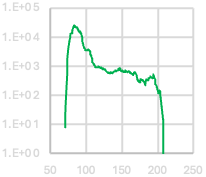

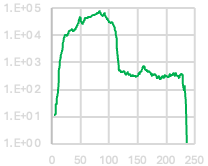

|  | 1824_fb |
| --- | --- |
| 0 | 0 |
| 1 | 0 |
| 2 | 0 |
| 3 | 0 |
| 4 | 0 |
| 5 | 0 |
| 6 | 0 |
| 7 | 0 |
| 8 | 0 |
| 9 | 0 |
| 10 | 0 |
| 11 | 0 |
| 12 | 0 |
| 13 | 0 |
| 14 | 0 |
| 15 | 0 |
| 16 | 0 |
| 17 | 0 |
| 18 | 0 |
| 19 | 0 |
| 20 | 0 |
| 21 | 0 |
| 22 | 0 |
| 23 | 0 |
| 24 | 0 |
| 25 | 0 |
| 26 | 0 |
| 27 | 0 |
| 28 | 0 |
| 29 | 0 |
| 30 | 0 |
| 31 | 0 |
| 32 | 0 |
| 33 | 0 |
| 34 | 0 |
| 35 | 0 |
| 36 | 0 |
| 37 | 0 |
| 38 | 0 |
| 39 | 0 |
| 40 | 0 |
| 41 | 0 |
| 42 | 0 |
| 43 | 0 |
| 44 | 0 |
| 45 | 0 |
| 46 | 0 |
| 47 | 0 |
| 48 | 0 |
| 49 | 0 |
| 50 | 0 |
| 51 | 0 |
| 52 | 0 |
| 53 | 0 |
| 54 | 0 |
| 55 | 0 |
| 56 | 0 |
| 57 | 0 |
| 58 | 0 |
| 59 | 0 |
| 60 | 0 |
| 61 | 0 |
| 62 | 0 |
| 63 | 0 |
| 64 | 0 |
| 65 | 0 |
| 66 | 0 |
| 67 | 0 |
| 68 | 0 |
| 69 | 0 |
| 70 | 8 |
| 71 | 25 |
| 72 | 148 |
| 73 | 642 |
| 74 | 1839 |
| 75 | 4025 |
| 76 | 6974 |
| 77 | 10109 |
| 78 | 12367 |
| 79 | 13211 |
| 80 | 14092 |
| 81 | 17628 |
| 82 | 21963 |
| 83 | 24758 |
| 84 | 24063 |
| 85 | 22524 |
| 86 | 22144 |
| 87 | 21957 |
| 88 | 20321 |
| 89 | 17546 |
| 90 | 15319 |
| 91 | 15444 |
| 92 | 14233 |
| 93 | 12635 |
| 94 | 8897 |
| 95 | 6941 |
| 96 | 5354 |
| 97 | 4300 |
| 98 | 3786 |
| 99 | 3548 |
| 100 | 3522 |
| 101 | 3612 |
| 102 | 3601 |
| 103 | 3826 |
| 104 | 3508 |
| 105 | 3005 |
| 106 | 2602 |
| 107 | 2260 |
| 108 | 1677 |
| 109 | 1349 |
| 110 | 1262 |
| 111 | 1100 |
| 112 | 1007 |
| 113 | 902 |
| 114 | 940 |
| 115 | 902 |
| 116 | 906 |
| 117 | 958 |
| 118 | 906 |
| 119 | 888 |
| 120 | 834 |
| 121 | 904 |
| 122 | 813 |
| 123 | 874 |
| 124 | 797 |
| 125 | 758 |
| 126 | 778 |
| 127 | 710 |
| 128 | 674 |
| 129 | 637 |
| 130 | 575 |
| 131 | 603 |
| 132 | 568 |
| 133 | 585 |
| 134 | 624 |
| 135 | 606 |
| 136 | 570 |
| 137 | 613 |
| 138 | 618 |
| 139 | 661 |
| 140 | 692 |
| 141 | 719 |
| 142 | 662 |
| 143 | 736 |
| 144 | 730 |
| 145 | 790 |
| 146 | 870 |
| 147 | 854 |
| 148 | 716 |
| 149 | 683 |
| 150 | 706 |
| 151 | 679 |
| 152 | 640 |
| 153 | 684 |
| 154 | 629 |
| 155 | 697 |
| 156 | 674 |
| 157 | 603 |
| 158 | 658 |
| 159 | 650 |
| 160 | 660 |
| 161 | 592 |
| 162 | 601 |
| 163 | 558 |
| 164 | 526 |
| 165 | 485 |
| 166 | 541 |
| 167 | 596 |
| 168 | 635 |
| 169 | 593 |
| 170 | 520 |
| 171 | 366 |
| 172 | 314 |
| 173 | 301 |
| 174 | 269 |
| 175 | 287 |
| 176 | 363 |
| 177 | 366 |
| 178 | 322 |
| 179 | 317 |
| 180 | 298 |
| 181 | 280 |
| 182 | 222 |
| 183 | 258 |
| 184 | 347 |
| 185 | 329 |
| 186 | 328 |
| 187 | 371 |
| 188 | 377 |
| 189 | 408 |
| 190 | 440 |
| 191 | 400 |
| 192 | 381 |
| 193 | 413 |
| 194 | 525 |
| 195 | 384 |
| 196 | 364 |
| 197 | 231 |
| 198 | 223 |
| 199 | 163 |

|  | IMG_1901.jpeg |
| --- | --- |
| 0 | 0 |
| 1 | 0 |
| 2 | 0 |
| 3 | 0 |
| 4 | 0 |
| 5 | 11 |
| 6 | 12 |
| 7 | 29 |
| 8 | 30 |
| 9 | 47 |
| 10 | 72 |
| 11 | 207 |
| 12 | 523 |
| 13 | 817 |
| 14 | 1129 |
| 15 | 1553 |
| 16 | 1854 |
| 17 | 2492 |
| 18 | 3742 |
| 19 | 5439 |
| 20 | 6887 |
| 21 | 8134 |
| 22 | 9312 |
| 23 | 10634 |
| 24 | 10240 |
| 25 | 10181 |
| 26 | 9759 |
| 27 | 10338 |
| 28 | 11509 |
| 29 | 13311 |
| 30 | 14236 |
| 31 | 14539 |
| 32 | 15524 |
| 33 | 15815 |
| 34 | 15753 |
| 35 | 16790 |
| 36 | 15964 |
| 37 | 14191 |
| 38 | 14407 |
| 39 | 15991 |
| 40 | 17188 |
| 41 | 17977 |
| 42 | 18880 |
| 43 | 20035 |
| 44 | 21758 |
| 45 | 25644 |
| 46 | 32365 |
| 47 | 38038 |
| 48 | 44932 |
| 49 | 49174 |
| 50 | 43129 |
| 51 | 34260 |
| 52 | 28200 |
| 53 | 27688 |
| 54 | 29875 |
| 55 | 32301 |
| 56 | 35297 |
| 57 | 38658 |
| 58 | 42941 |
| 59 | 45031 |
| 60 | 44010 |
| 61 | 45773 |
| 62 | 45704 |
| 63 | 46385 |
| 64 | 46847 |
| 65 | 48441 |
| 66 | 46764 |
| 67 | 47666 |
| 68 | 49371 |
| 69 | 51990 |
| 70 | 53010 |
| 71 | 56138 |
| 72 | 59378 |
| 73 | 58180 |
| 74 | 56386 |
| 75 | 58389 |
| 76 | 62471 |
| 77 | 62003 |
| 78 | 61743 |
| 79 | 65252 |
| 80 | 67121 |
| 81 | 67867 |
| 82 | 73297 |
| 83 | 79662 |
| 84 | 74232 |
| 85 | 69433 |
| 86 | 62339 |
| 87 | 56661 |
| 88 | 50655 |
| 89 | 50912 |
| 90 | 56510 |
| 91 | 64772 |
| 92 | 65504 |
| 93 | 58654 |
| 94 | 48074 |
| 95 | 42899 |
| 96 | 39780 |
| 97 | 41075 |
| 98 | 36203 |
| 99 | 33800 |
| 100 | 31956 |
| 101 | 31135 |
| 102 | 29715 |
| 103 | 29454 |
| 104 | 30648 |
| 105 | 31346 |
| 106 | 29571 |
| 107 | 26523 |
| 108 | 23021 |
| 109 | 18877 |
| 110 | 16270 |
| 111 | 13283 |
| 112 | 10661 |
| 113 | 7473 |
| 114 | 3549 |
| 115 | 1417 |
| 116 | 696 |
| 117 | 523 |
| 118 | 486 |
| 119 | 541 |
| 120 | 483 |
| 121 | 460 |
| 122 | 452 |
| 123 | 436 |
| 124 | 458 |
| 125 | 490 |
| 126 | 470 |
| 127 | 461 |
| 128 | 398 |
| 129 | 353 |
| 130 | 400 |
| 131 | 361 |
| 132 | 434 |
| 133 | 396 |
| 134 | 358 |
| 135 | 357 |
| 136 | 376 |
| 137 | 348 |
| 138 | 313 |
| 139 | 325 |
| 140 | 317 |
| 141 | 337 |
| 142 | 304 |
| 143 | 313 |
| 144 | 320 |
| 145 | 323 |
| 146 | 305 |
| 147 | 282 |
| 148 | 324 |
| 149 | 310 |
| 150 | 323 |
| 151 | 393 |
| 152 | 368 |
| 153 | 381 |
| 154 | 446 |
| 155 | 433 |
| 156 | 461 |
| 157 | 515 |
| 158 | 680 |
| 159 | 691 |
| 160 | 721 |
| 161 | 703 |
| 162 | 685 |
| 163 | 584 |
| 164 | 526 |
| 165 | 506 |
| 166 | 493 |
| 167 | 473 |
| 168 | 441 |
| 169 | 479 |
| 170 | 465 |
| 171 | 429 |
| 172 | 426 |
| 173 | 380 |
| 174 | 373 |
| 175 | 314 |
| 176 | 354 |
| 177 | 365 |
| 178 | 338 |
| 179 | 371 |
| 180 | 368 |
| 181 | 339 |
| 182 | 356 |
| 183 | 299 |
| 184 | 273 |
| 185 | 275 |
| 186 | 311 |
| 187 | 312 |
| 188 | 269 |
| 189 | 243 |
| 190 | 239 |
| 191 | 254 |
| 192 | 287 |
| 193 | 309 |
| 194 | 288 |
| 195 | 301 |
| 196 | 278 |
| 197 | 293 |
| 198 | 331 |
| 199 | 317 |

C	D

g	g

Fig. S2. Fluorescence images used in lipid deposition in embryos (A) and lipofuscin accumulation analysis (B-D). A: Overlay of the flight field and Nile Red fluorescence images taken of each neonate. B: An example of adult female nephridium image. C, D: Example of adult female mid-body and abdomen, respectively, images with lumps of lipofuscins visible (green). g = gut. Inserts: histograms of green channel fluorescence, intensities above 100 corresponding to lipofuscin lumps.

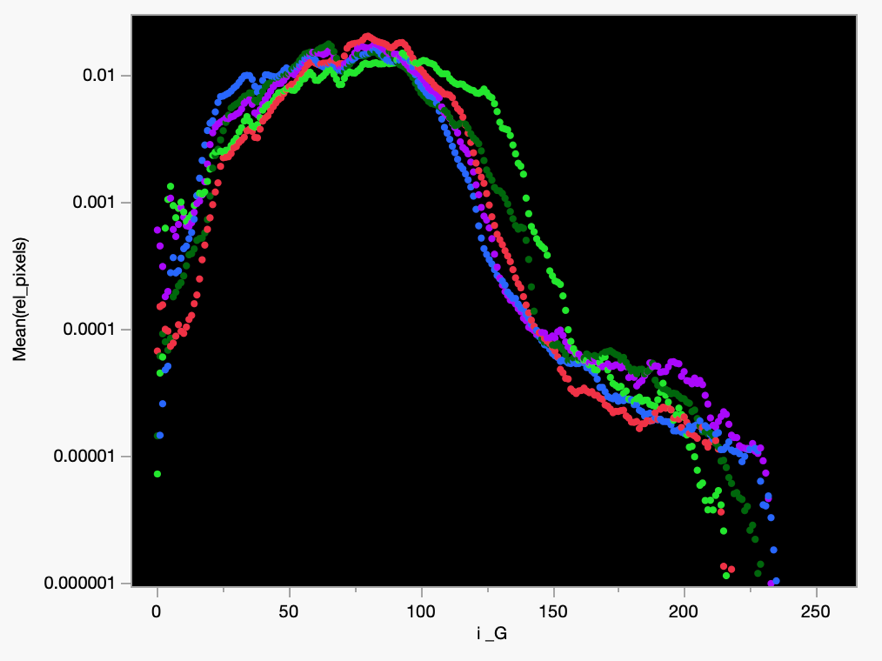

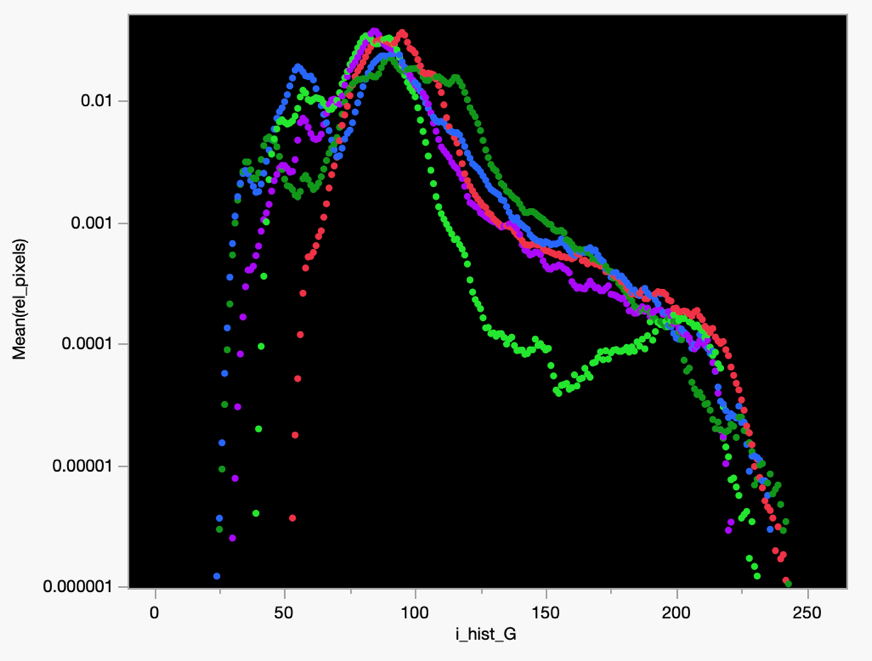
A: abdominal fat body	B: thoracal fat body

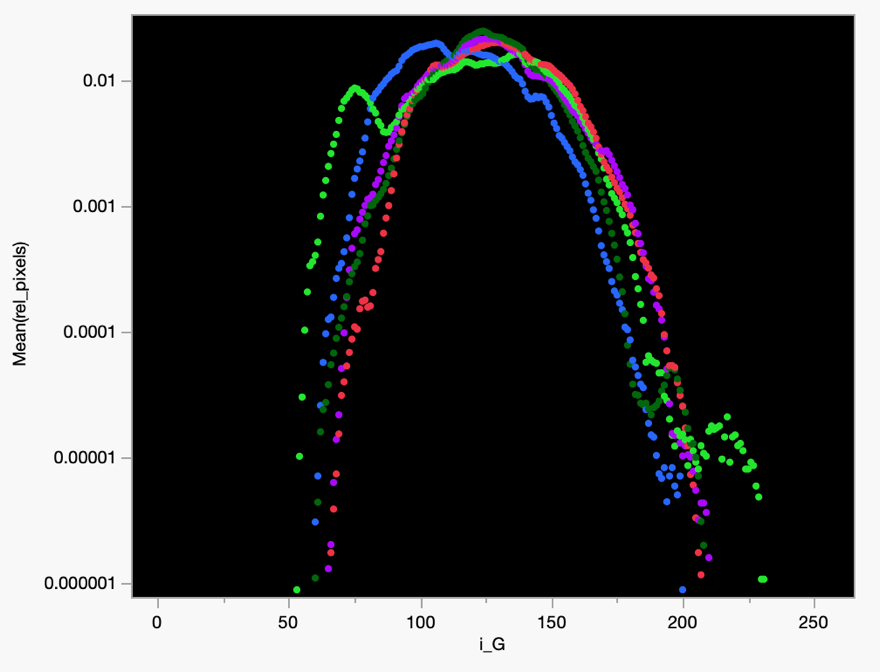

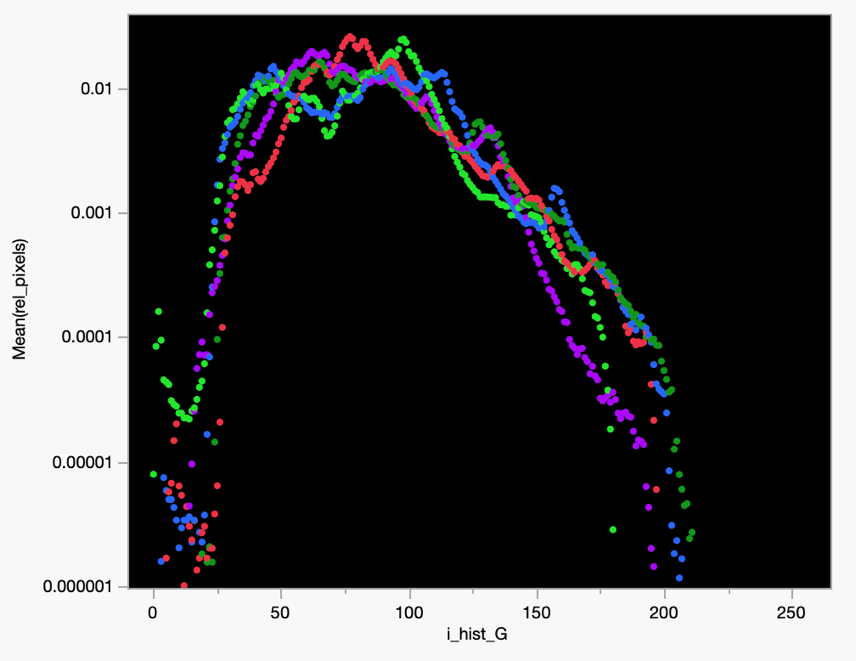
C: ovary	D: nephridium

Fig. S2.3new. Green channel fluorescence intensity profiles in 4 ROIs in *Daphnia* exposed to 5 different treatments: high and low food control (dark and light green), BHB (red), NMN (blue), and both BHB and NMN (purple). Data for each intensity averaged across replicates.

Results

Fig. S3. Autofluorescence histograms in 4 RoIs. Dark and light green: full diet and CR diet, respectively; red: BHB treatment; blue: NMN treatment; purple: combined BHB and NMN treatment.

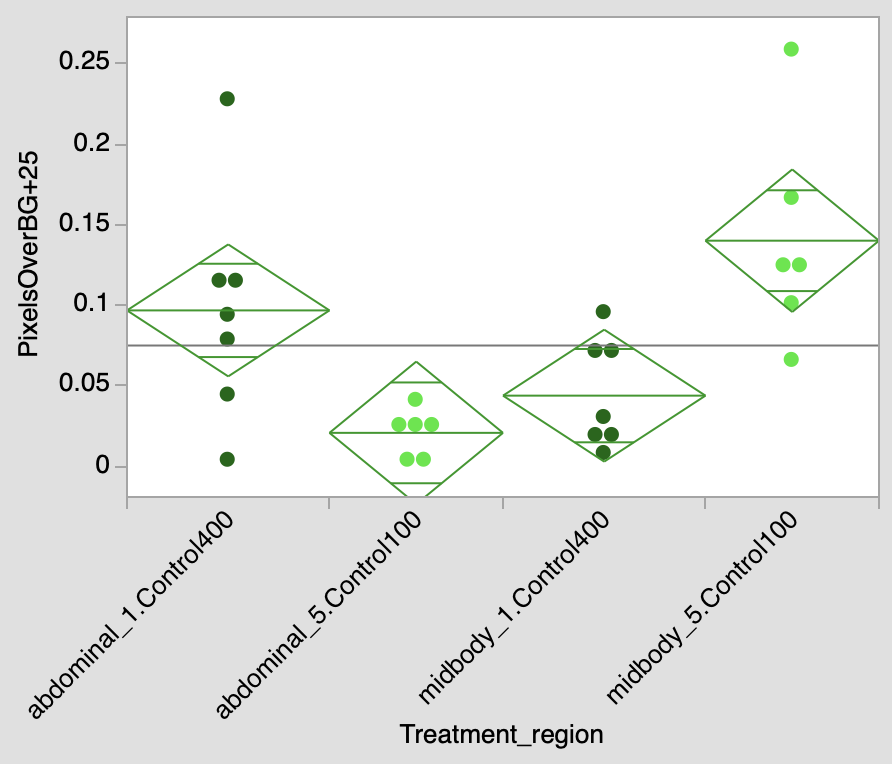

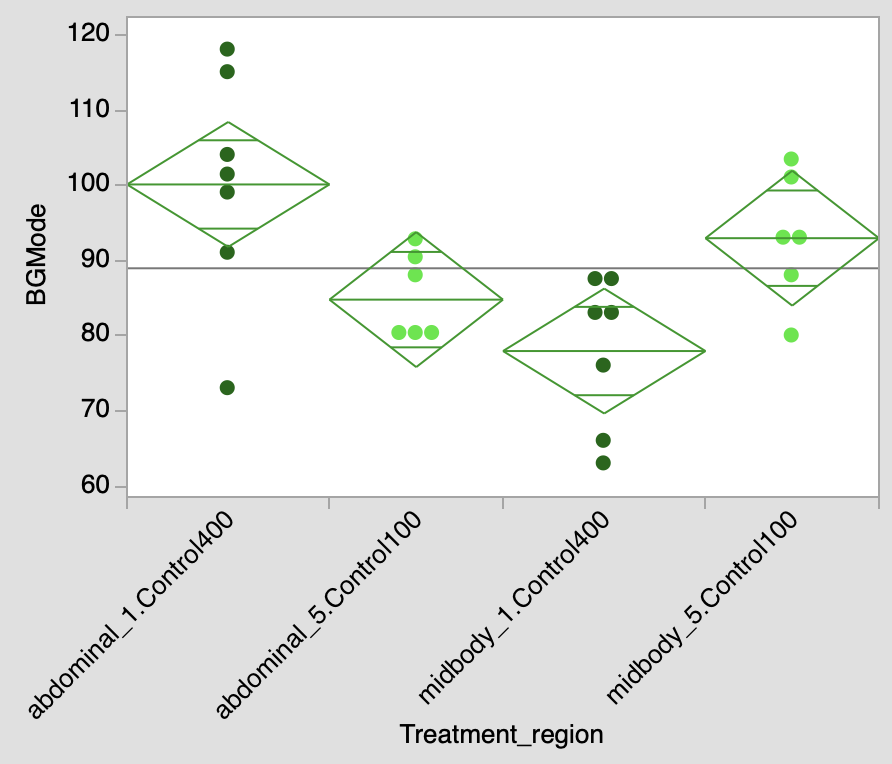
A	B

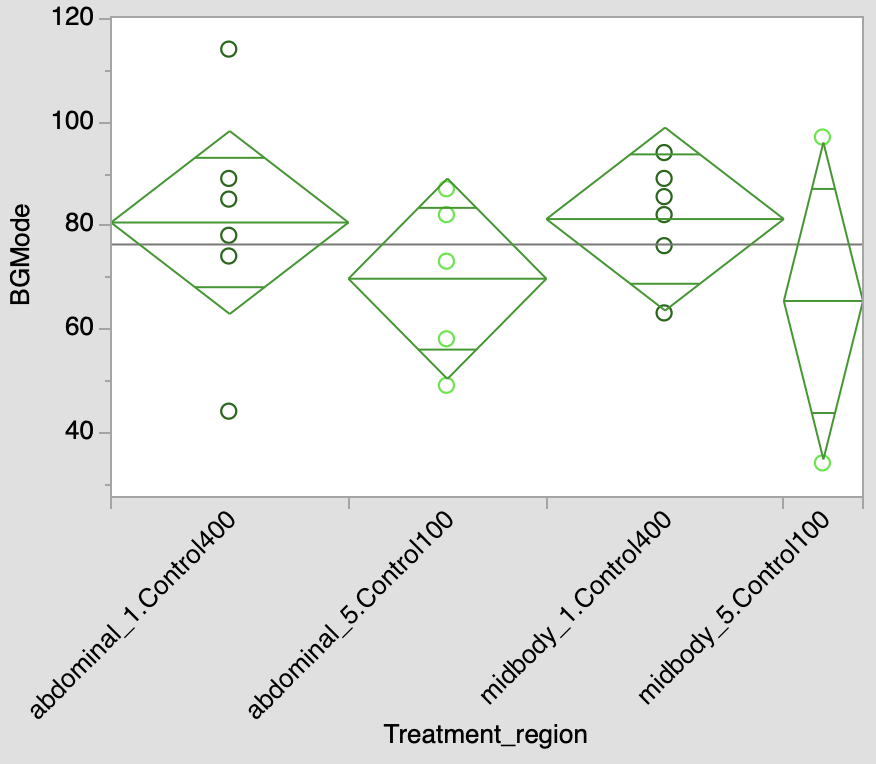

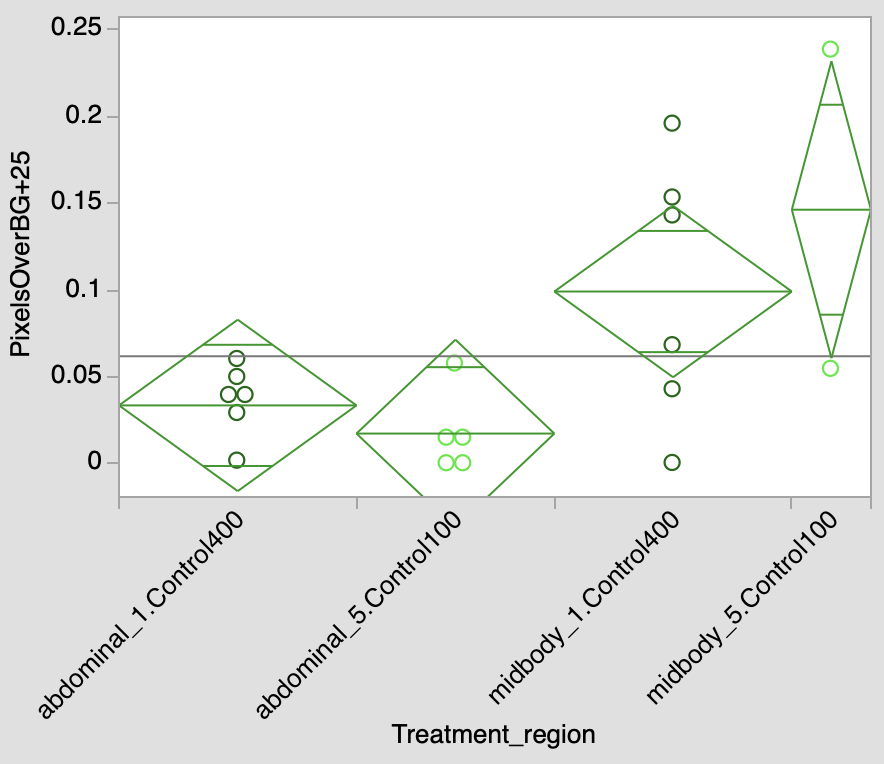
C	D

Fig. S4. Food level (4E5 vs 1E5 cells/mL/day) X body region (abdomen vs. midbody) interaction in fat body background fluorescence (A) and portion of fluorescence above background + 25 units (B). Inserts: 2-way ANOVA.

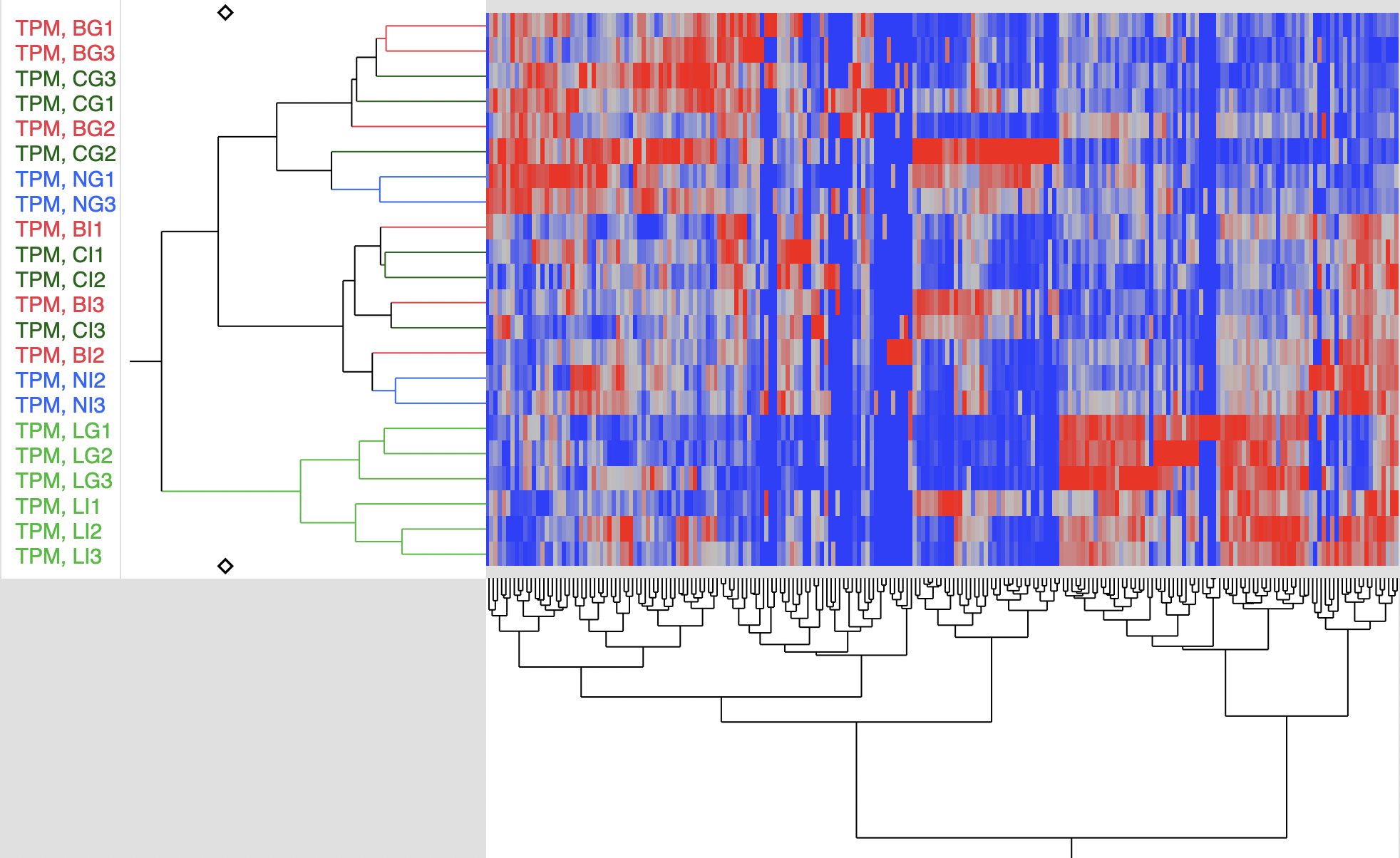

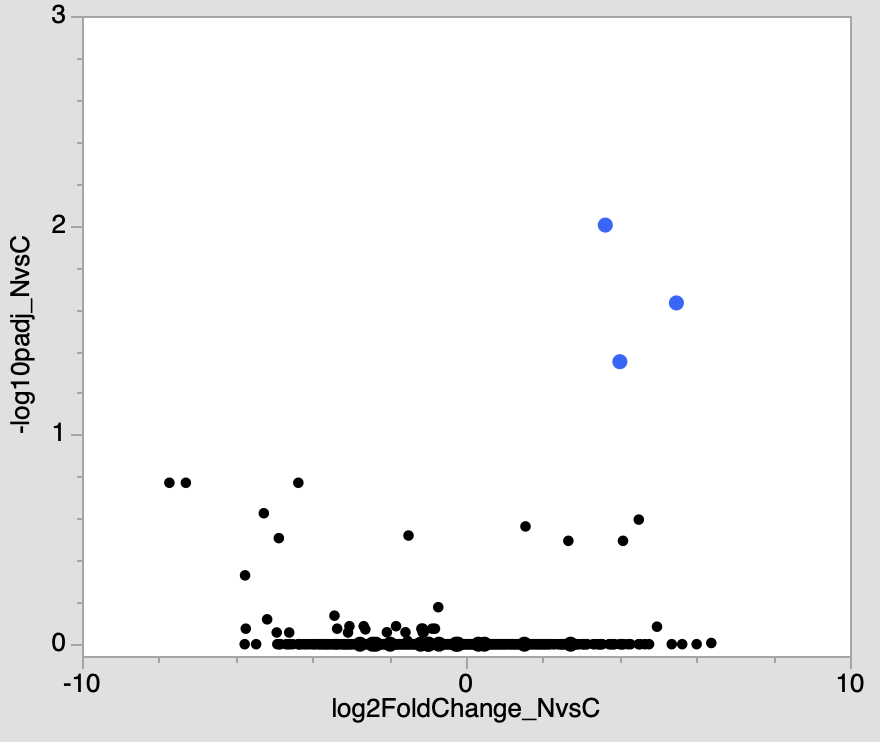

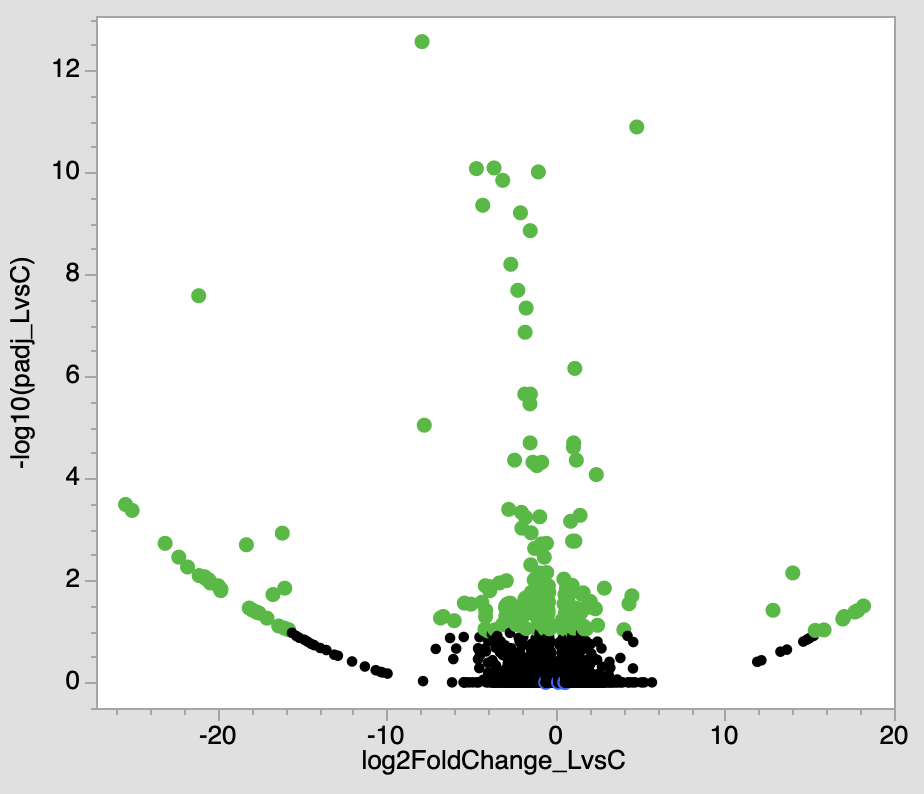
B	C	D

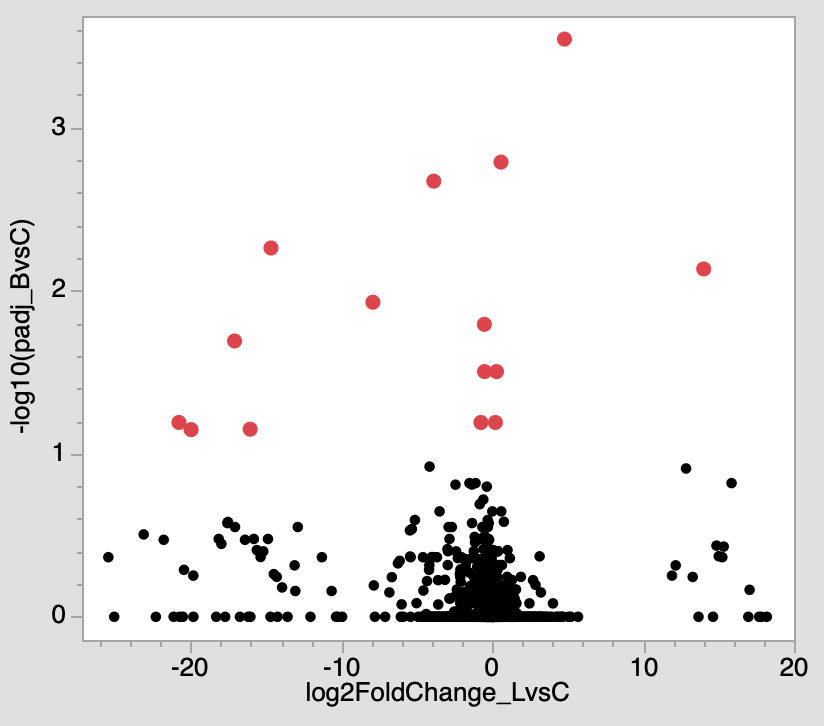

Fig S5. A. Two-way clustering heatmap of genes up (red) and down (blue) regulated in CR treatment (green labels), BHB treatment (red labels), and NMN treatment (blue labels). “I” and “G” letters in the labels indicate clones (IL and GB). B – C: volcano plots for DE for CR (B), BHB (C), and NMN (D) effects.

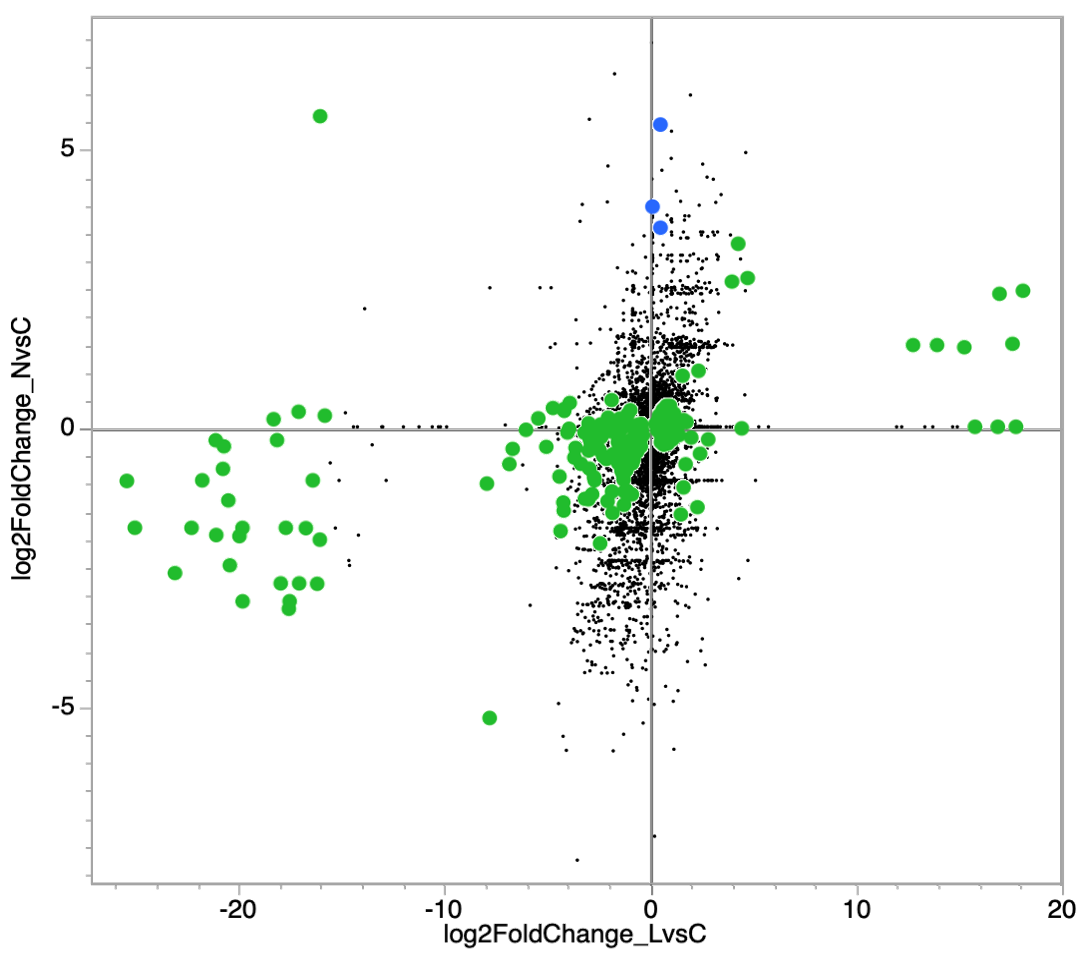

Log_2_(FoldChange) full vs. CR

Log_2_(FoldChange) full vs. NMN

Fig. S6. “2-way” volcano plot for the effects of CR and NMN exposure (cf. Main text Fig. 5 B for the BHB exposure). Green: genes with a significant DE in response to CR. Blue: genes with a significant DE in response to NMN. Genes with neither shown as black dots.

Table S1. ANOVAs of the effects of treatments and clones on green channel fluorescence from tissues background and from lipofuscin lumps. P-values < 0.002 shown in bold. See main text. Fig. X for Dunnett post-hoc comparisons of each treatment to 4E5 cells/ml/day control, where significant effects are indicated here.

| ROI = abdominal fat body | | | Response: pixels over 100  (background) | | |  | | Response: pixels over 25, background-subtracted  (lipofuscin lumps) | | |
| --- | --- | --- | --- | --- | --- | --- | --- | --- | --- | --- |
| Source | DF | Sum of Squares | F Ratio | Prob > F |  | | Sum of Squares | | F Ratio | Prob > F |
| Treatments | 4 | 0.726 | 3.755 | **0.01** |  | | 0.022 | | 1.407 | 0.25 |
| Clone | 1 | 0.515 | 10.666 | **0.002** |  | | 0.012 | | 3.199 | 0.08 |
| Treatments*Clone | 4 | 0.315 | 1.63 | 0.18 |  | | 0.012 | | 0.78 | 0.55 |
| Error | 50 | 2.415 |  |  |  | | 0.192 | |  |  |
| ROI = thoracal fat body | | |  |  |  | |  | |  |  |
| Source | DF | Sum of Squares | F Ratio | Prob > F |  | | Sum of Squares | | F Ratio | Prob > F |
| Treatments | 4 | 0.101 | 2.809 | 0.04 |  | | 0.04 | | 1.483 | 0.22 |
| Clone | 1 | 0.016 | 1.815 | 0.18 |  | | 0.012 | | 1.827 | 0.18 |
| Treatments*Clone | 4 | 0.111 | 3.08 | 0.03 |  | | 0.003 | | 0.13 | 0.97 |
| Error | 44 | 0.397 |  |  |  | | 0.299 | |  |  |
| ROI = nephridium | | |  |  |  | |  | |  |  |
| Source | DF | Sum of Squares | F Ratio | Prob > F |  | | Sum of Squares | | F Ratio | Prob > F |
| Treatments | 4 | 0.439 | 3.077 | 0.025 |  | | 0.074 | | 1.506 | 0.21 |
| Clone | 1 | 0.163 | 4.588 | 0.04 |  | | 0.017 | | 1.422 | 0.24 |
| Treatments*Clone | 4 | 0.375 | 2.63 | 0.05 |  | | 0.077 | | 1.58 | 0.19 |
| Error | 50 | 1.781 |  |  |  | | 0.613 | |  |  |
| ROI = ovary | | |  |  |  | |  | |  |  |
| Source | DF | Sum of Squares | F Ratio | Prob > F |  | | Sum of Squares | | F Ratio | Prob > F |
| Treatments | 4 | 0.092 | 0.338 | 0.851 |  | | 0.033 | | 0.607 | 0.66 |
| Clone | 1 | 0.799 | 11.746 | **0.0012** |  | | 0.181 | | 13.095 | **0.0007** |
| Treatments*Clone | 4 | 0.247 | 0.91 | 0.47 |  | | 0.028 | | 0.51 | 0.73 |
| Error | 50 | 3.403 |  |  |  | | 0.69 | |  |  |

Table S2. 3-way ANOVAs of the differences between food levels (1E5 vs. 4E5 cells/mL/day), clones, and body regions (thoracal fat body vs. abdominal fat body) in background green autofluorescence (portions of pixels above 100 units, left) and background-subtracted autofluorescence from highly fluorescent areas (lipofuscin lumps). P-values < 0.002 shown in bold.

| ROI = abdominal vs. thoracal fat body | | | Response: pixels over 100  (background) | | | |  | | Response: pixels over 25, background-subtracted  (lipofuscin lumps) | | |
| --- | --- | --- | --- | --- | --- | --- | --- | --- | --- | --- | --- |
| Source | DF | Sum of Squares | | F Ratio | Prob > F |  | | Sum of Squares | | F Ratio | Prob > F |
| Treatments | 1 | 0.115 | | 3.256 | 0.08 |  | | 0.002 | | 0.539 | 0.47 |
| Clone | 1 | 0.147 | | 4.156 | 0.05 |  | | 0.00001 | | 0.004 | 0.95 |
| Treatments*Clone | 1 | 0.002 | | 0.066 | 0.8 |  | | 0.00007 | | 0.024 | 0.88 |
| ROI | 1 | 0.01 | | 0.27 | 0.61 |  | | 0.041 | | 14.054 | **0.001** |
| Treatments*ROI | 1 | 0.473 | | 13.405 | **0.001** |  | | 0.034 | | 11.47 | **0.002** |
| Clone*ROI | 1 | 0.054 | | 1.524 | 0.22 |  | | 0.01 | | 3.384 | 0.07 |
| Treatments*Clone*ROI | 1 | 0.287 | | 8.126 | **0.007** |  | | 0.007 | | 2.455 | 0.126 |
| Error | 37 | 1.305 | |  |  |  | | 0.108 | |  |  |

Table S3. Concordance between genes with significant DE in response to CR and in response to BHB or NMN exposure, relative to the full diet control. Criteria of significant DE: p_adj_<0.1 (top) and p_adj_<0.1 & |log2(FC)|>1 (bottom). P-values are from a 2-tailed Fisher Exact test.

| Criterion of significance | |  |  |  |
| --- | --- | --- | --- | --- |
| p_adj_<0.1 |  |  | BHB vs full diet control | NMN vs full diet control |
|  | Number of genes with DE |  | 15 | 4 |
| CR vs full diet control | 216 | Expected | 0.392 | 0.1047 |
|  |  | Observed | 9 | 0 |
|  |  | FET P-value | <.0001 | 1 |
| BHB vs full diet control | 15 | Expected | - | 0.0067 |
|  |  | Observed | - | 0 |
|  |  | FET P-value | - | 1 |
| p_adj_<0.1 & \|log2(FC)\|>1 | |  | B vs C | L vs C |
|  | Number of genes with DE |  | 14 | 4 |
| CR vs full diet control | 141 | Expected | 0.239 | 0.068 |
|  |  | Observed | 8 | 0 |
|  |  | FET P-value | <.0001 | 1 |
| BHB vs full diet control | 14 | Expected | - | 0.0063 |
|  |  | Observed | - | 0 |
|  |  | FET P-value | - | 1 |

Table S4. Genes up- and downregulated in response to CR, BHB, or NMN treatments with p_adj_<0.1 and |log_2_FC|>1, relative to full diet control. Assembly ID shown when UniProt protein ID not available

| UniProtKB | | Description | | PANTHERDescription | log_2_FC | padj | |
| --- | --- | --- | --- | --- | --- | --- | --- |
| Up in CR | |  | |  |  |  | |
| KZR96933 | | gag-pol polyprotein precursor | | GUANINE NUCLEOTIDE-BINDING PROTEIN G(I)_G(S)_G(O) SUBUNIT GAMMA-2 ISOFORM X1-RELATED | 18.198 | 0.032 | |
| KZR96382 | | Uncharacterized protein | | RIBONUCLEASE H : PHD-TYPE DOMAIN-CONTAINING PROTEIN | 17.855 | 0.039 | |
| KZS05075 | | Apoptosis 1 inhibitor | | DEATH-ASSOCIATED INHIBITOR OF APOPTOSIS 2 | 17.691 | 0.042 | |
| KZS01839 | | polyprotein of retroviral origin | |  | 17.056 | 0.051 | |
| KZR98917 | | Structural maintenance of chromosomes protein 2 | | | 17.001 | 0.057 | |
| maker-falcon_000039F-snap-gene-0.22 | | Uncharacterized protein | |  | 16.972 | 0.057 | |
| KZS15433 | | Hepatoma-derived growth factor-related protein 3 | | OXIDOREDUCTASE GLYR1-RELATED | 15.862 | 0.094 | |
| KZS03157 | | Uncharacterized protein | | CXC DOMAIN-CONTAINING PROTEIN | 15.341 | 0.096 | |
| KZS02589 | | Uncharacterized protein | | 60S RIBOSOMAL PROTEIN L31 | 14.014 | 0.007 | |
| maker-falcon_000004F-snap-gene-36.11 | | Juvenile hormone epoxide hydrolase 1, Juvenile hormone epoxide hydrolase I, RH03631p | | | 12.846 | 0.039 | |
| KZS14798 | | Zinc metalloproteinase nas-13 | | DISCOIDIN, CUB, EGF, LAMININ , AND ZINC METALLOPROTEASE DOMAIN CONTAINING | 4.789 | <0.001 | |
| KZS13722 | | Uncharacterized protein | | FIBROUS SHEATH CABYR-BINDING PROTEIN | 4.501 | 0.02 | |
| KZS03691 | | pol protein | | GH03217P-RELATED | 4.327 | 0.029 | |
| KZS02262 | | Uncharacterized protein | |  | 4.025 | 0.093 | |
| KZS08369 | | Acetylcholinesterase collagenic tail peptide | | ENDOSTATIN DOMAIN-CONTAINING PROTEIN | 2.869 | 0.014 | |
| KZS13186 | | Ferric-chelate reductase 1,EC=1.-.-.-/sw | | CYTOCHROME B561/FERRIC REDUCTASE TRANSMEMBRANE | 2.473 | 0.076 | |
| KZS18390 | | Puromycin-sensitive aminopeptidase protein | | AMINOPEPTIDASE | 2.396 | <0.001 | |
| KZS13750 | | Uncharacterized protein | | SUBFAMILY NOT NAMED | 2.344 | 0.036 | |
| KZS08312 | | Phospholipase A2 | | PHOSPHOLIPASE A2 | 2.043 | 0.026 | |
| maker-falcon_000000F-snap-gene-71.71 | | Uncharacterized protein | |  | 1.809 | 0.089 | |
| KZS13988 | | Solute carrier family 22 member 7 | | BETA-ALANINE TRANSPORTER | 1.768 | 0.035 | |
| KZS16840 | | Uncharacterized protein | |  | 1.677 | 0.089 | |
| KZS14007 | | Uncharacterized protein | | CHLOROPHYLLASE-2 | 1.666 | 0.079 | |
| maker-falcon_000001F-snap-gene-42.78 | | Uncharacterized protein | |  | 1.618 | 0.018 | |
| KZS05952 | | Bone morphogenetic protein | | IM:7138239 | 1.539 | 0.019 | |
| KZS12089 | | Middleman of seventy-eight signaling | | STERILE ALPHA MOTIF DOMAIN-CONTAINING PROTEIN 7 ISOFORM X1 | 1.469 | 0.043 | |
| KZS15794 | | Uncharacterized protein | | OS09G0480532 PROTEIN | 1.436 | 0.001 | |
| KZS03382 | | Uncharacterized protein | |  | 1.276 | 0.055 | |
| KZS05953 | | Bone morphogenetic protein 10 | | TGF_BETA_2 DOMAIN-CONTAINING PROTEIN | 1.214 | <0.001 | |
| KZS03894 | | Uncharacterized protein | | HOMEOTIC PROTEIN SEX COMBS REDUCED | 1.204 | 0.035 | |
| KZS13708 | | Clip-domain serine protease | | FI16631P1-RELATED | 1.129 | 0.002 | |
| KZS15950 | | Alanine--tRNA ligase, cytoplasmic | | ASPARTATE DEHYDROGENASE DOMAIN-CONTAINING PROTEIN : L-ASPARTATE DEHYDROGENASE-RELATED | 1.125 | <0.001 | |
| KZS13283 | | Uncharacterized protein | |  | 1.062 | 0.087 | |
| KZS11268 | | Bric-a-brac | | RIBBON, ISOFORM C | 1.056 | <0.001 | |
| KZS21204 | | Uncharacterized protein | |  | 1.045 | <0.001 | |
| KZS03019 | | ATP-binding cassette sub-family G member | | FI03229P | 1.025 | 0.074 | |
| KZS14500 | | Tetraspanin 68C | | TETRASPANIN | 1.006 | 0.035 | |
| Down in CR | |  | |  |  |  | |
| KZS20257 | | Uncharacterized protein | | GUANINE NUCLEOTIDE-BINDING PROTEIN G(I)_G(S)_G(O) SUBUNIT GAMMA-2 ISOFORM X1-RELATED | -25.444 | <0.001 | |
| KZS16882 | | Uncharacterized protein | |  | -25.054 | <0.001 | |
| KZS04169 | | TE: Polyprotein of retroviral origin | | SUBFAMILY NOT NAMED | -23.102 | 0.002 | |
| KZS10178 | | TE: Retrovirus-related Pol polyprotein from transposon | | RIBONUCLEASE H | -22.295 | 0.004 | |
| KZS02879 | | Uncharacterized protein | |  | -21.774 | 0.005 | |
| KZS06880 | | pupal cuticle protein | | CUTICULAR PROTEIN 49AA-RELATED | -21.117 | <0.001 | |
| KZR97400 | | Uncharacterized protein | |  | -21.091 | 0.008 | |
| snap_masked-falcon_000001F-processed-gene-41.107 | | Chitin deacetylase 9 | |  | -20.765 | 0.009 | |
| maker-falcon_000112F-augustus-gene-0.75 | | LambdaTry | |  | -20.718 | 0.009 | |
| KZS20343 | | Uncharacterized protein | | THAP DOMAIN CONTAINING 9 | -20.503 | 0.01 | |
| KZS07568 | | Uncharacterized protein | |  | -20.43 | 0.011 | |
| KZS05290 | | cuticle protein | | CUTICULAR PROTEIN 50CB-RELATED | -19.962 | 0.013 | |
| KZR98961 | | Uncharacterized protein | |  | -19.813 | 0.016 | |
| maker-falcon_000010F-snap-gene-23.55 | | Carboxyl/cholinesterase | |  | -19.806 | 0.015 | |
| snap_masked-falcon_000023F-processed-gene-11.66 | | | |  | -18.296 | 0.002 | |
| maker-falcon_000030F-augustus-gene-4.87 | | Uncharacterized protein | |  | -18.128 | 0.035 | |
| KZS07453 | | Uncharacterized protein daphplx:hxJGI_V11_99747 | | TY3_CAPSID DOMAIN-CONTAINING PROTEIN | -17.951 | 0.038 | |
| KZR98257 | | Uncharacterized protein | |  | -17.701 | 0.042 | |
| KZS00071 | | Zinc finger protein | |  | -17.655 | 0.042 | |
| KZS13114 | | Uncharacterized protein | |  | -17.654 | 0.042 | |
| KZS17402 | | Uncharacterized protein | |  | -17.554 | 0.044 | |
| KZS03769 | | Uncharacterized protein ENSANGP00000027469 | | REVERSE TRANSCRIPTASE | -17.516 | 0.045 | |
| KZS20869 | | ANK, Ankyrin repeats | | TRANSPOSASE | -17.077 | 0.055 | |
| KZR96704 | | Uncharacterized protein | |  | -17.051 | 0.055 | |
| KZS04909 | | Uncharacterized protein | |  | -16.725 | 0.019 | |
| KZS17095 | | Uncharacterized protein | |  | -16.387 | 0.078 | |
| KZS04928 | | Uncharacterized protein | | TRANSPOSASE | -16.173 | 0.001 | |
| KZS03747 | | Uncharacterized protein | | BEN DOMAIN-CONTAINING PROTEIN 6 | -16.043 | 0.087 | |
| KZS21741 | | Uncharacterized protein | | AT07410P | -16.028 | 0.014 | |
| KZS21811 | | Uncharacterized protein | | SUBFAMILY NOT NAMED | -15.8 | 0.093 | |
| KZS15417 | | Uncharacterized protein | | PROTEIN, PUTATIVE-RELATED | -7.911 | <0.001 | |
| KZS08918 | | Clip-domain serine protease | | CLIP-DOMAIN SERINE PROTEASE | -7.778 | <0.001 | |
| KZS10251 | | TE: Polyprotein of retroviral origin | | RIBONUCLEASE H | -6.817 | 0.055 | |
| KZS06862 | | ppn | | KUNITZ-TYPE PROTEASE INHIBITOR-RELATED | -6.656 | 0.051 | |
| KZS17935 | | Uncharacterized protein | |  | -6.007 | 0.062 | |
| KZS10750 | | Cuticular protein 49Ag | | CUTICULAR PROTEIN 49AA-RELATED | -5.408 | 0.028 | |
| KZS12842 | | Uncharacterized protein | |  | -5.018 | 0.029 | |
| genemark-falcon_000004F-processed-gene-32.11 | | | |  | -4.694 | <0.001 | |
| KZS06881 | | Uncharacterized protein | | CUTICULAR PROTEIN 49AA-RELATED | -4.376 | 0.027 | |
| KZS15419 | | Uncharacterized protein | |  | -4.321 | <0.001 | |
| KZR96189 | | Pol protein | |  | -4.19 | 0.09 | |
| KZS12566 | | Uncharacterized protein | | SI:CH211-282J17.13-RELATED | -4.169 | 0.013 | |
| KZS18515 | | cuticle protein | | CUTICULAR PROTEIN 50CB-RELATED | -4.154 | 0.051 | |
| KZS04118 | | Uncharacterized protein | |  | -4.144 | 0.039 | |
| KZS12785 | | Uncharacterized protein | | ZGC:174862 | -3.985 | 0.098 | |
| KZS16709 | | Uncharacterized protein | | INSULIN-LIKE 3 | -3.908 | 0.016 | |
| KZS03586 | | Uncharacterized protein | |  | -3.875 | 0.013 | |
| KZS16639 | | Gastric triacylglycerol lipase precursor | | LIPASE-RELATED | -3.652 | <0.001 | |
| KZS17431 | | Aristaless-related homeobox protein | | CUTICULAR PROTEIN 50CB-RELATED | -3.594 | 0.093 | |
| KZR95740 | | Uncharacterized protein | |  | -3.333 | 0.011 | |
| KZS16126 | | Uncharacterized protein | |  | -3.152 | 0.074 | |
| KZS13187 | | Defense protein l(2)34Fc | | DEFENSE PROTEIN L(2)34FC | -3.137 | <0.001 | |
| KZS03189 | | Copper-zinc cu-zn superoxide dismutase. | | VACUOLAR PROTEIN SORTING-ASSOCIATED PROTEIN 33A | -2.968 | 0.034 | |
| KZS05677 | | TE: Retrovirus-related Pol polyprotein from transposon TNT 1-94 | | TRANSPOSON TY4-P GAG-POL POLYPROTEIN | -2.957 | 0.051 | |
| KZS06786 | | Uncharacterized protein | | GLYCINE-RICH CELL WALL STRUCTURAL PROTEIN 2 | -2.948 | 0.031 | |
| KZS14969 | | Uncharacterized protein | |  | -2.926 | 0.01 | |
| KZS06888 | | pupal cuticle protein | | CUTICULAR PROTEIN 49AA-RELATED | -2.891 | 0.054 | |
| KZS06888 | | pupal cuticle protein | | CUTICULAR PROTEIN 49AA-RELATED | -2.84 | 0.029 | |
| KZS17536 | | Pro-resilin | | CUTICULAR PROTEIN 50CB-RELATED | -2.798 | 0.074 | |
| KZS12430 | | Uncharacterized protein | | AGAP007094-PA | -2.785 | <0.001 | |
| KZS12571 | | Glioma pathogenesis-related protein | | AT27079P-RELATED | -2.692 | 0.028 | |
| KZS19569 | | Serine proteinase | | BCDNA.GH08420-RELATED | -2.662 | <0.001 | |
| KZS12501 | | Sex-determining protein fem-1 | | PROTEIN FEM-1 HOMOLOG CG6966 | -2.437 | <0.001 | |
| KZS17717 | | G-protein-coupled receptor | | DOPAMINE RECEPTOR 2 | -2.397 | 0.035 | |
| maker-falcon_000066F-snap-gene-3.144 | | neurexin IV | |  | -2.374 | 0.04 | |
| KZS05838 | | Uncharacterized protein | | ZGC:194578 | -2.343 | 0.074 | |
| KZS18148 | | FAM162B | | UPF0389 PROTEIN CG9231 | -2.245 | <0.001 | |
| KZS11900 | | Uncharacterized protein | |  | -2.09 | <0.001 | |
| KZS16109 | | Bis(5'-adenosyl)-triphosphatase | | BIS(5'-ADENOSYL)-TRIPHOSPHATASE | -2.031 | <0.001 | |
| KZS14750 | | Chitin deacetylase 9 | | AQUARIUS-RELATED | -2.005 | 0.001 | |
| KZS14744 | | clip-domain serine protease | | AQUARIUS-RELATED | -1.868 | 0.047 | |
| KZS21012 | | Vitellogenin-1 precursor | | VITELLOGENIN-1-RELATED | -1.835 | 0.024 | |
| KZS13674 | | Uncharacterized protein | | GLYCOPROTEIN LIG-1, PUTATIVE-RELATED | -1.834 | <0.001 | |
| KZS03251 | | Cathepsin E-A | |  | -1.814 | <0.001 | |
| KZS13699 | | Chitinase | | CHITINASE | -1.79 | 0.022 | |
| KZS19327 | | Uncharacterized protein | | PROTEIN ROADKILL | -1.789 | 0.001 | |
| KZS11460 | | Ribonuclease P protein subunit p40 | | C1Q DOMAIN-CONTAINING PROTEIN | -1.758 | 0.088 | |
| KZS05919 | | Neuronal growth regulator 1 | | DPR-INTERACTING PROTEIN ETA, ISOFORM B-RELATED | -1.754 | <0.001 | |
| KZS21034 | | UPF0160 protein MYG1, mitochondrial | | MYG1 EXONUCLEASE | -1.735 | 0.043 | |
| KZS18583 | | ATP-binding cassette sub-family B member | | ATP-DEPENDENT TRANSLOCASE ABCB1 | -1.528 | <0.001 | |
| KZS10136 | | Uncharacterized protein | | 1,4-BETA-D-GLUCAN CELLOBIOHYDROLASE B | -1.518 | <0.001 | |
| KZS08085 | | Nose resistant to fluoxetine protein | | O-ACYLTRANSFERASE | -1.511 | <0.001 | |
| KZR98997 | | Uncharacterized protein | | BTB DOMAIN-CONTAINING PROTEIN | -1.503 | <0.001 | |
| KZS13265 | | tartan | | GLYCOPROTEIN LIG-1, PUTATIVE-RELATED | -1.481 | 0.005 | |
| KZS16921 | | Uncharacterized protein | | GANGLIOSIDE GM2 ACTIVATOR : GM2 GANGLIOSIDE ACTIVATOR PROTEIN | -1.457 | 0.001 | |
| KZS16666 | | Sulfotransferase sult | | PROTEIN, PUTATIVE-RELATED : TRANSFERASE, PUTATIVE-RELATED | -1.44 | 0.017 | |
| KZS11831 | | Brain chitinase and chia | |  | -1.361 | 0.043 | |
| KZR97438 | | C-type lectin domain family 10 member A | | AT17652P-RELATED | -1.352 | <0.001 | |
| KZS10092 | | btb/poz domain-containing protein | |  | -1.319 | 0.019 | |
| KZS13207 | | Chitin deacetylase 4 | | CHITIN DEACETYLASE-LIKE 9, ISOFORM A | -1.308 | 0.04 | |
| KZS03319 | | Carboxypeptidase B | | PEPTIDASE M14 CARBOXYPEPTIDASE A DOMAIN-CONTAINING PROTEIN | -1.277 | 0.01 | |
| KZS18537 | | Uncharacterized protein | | AGAP002848-PA | -1.254 | 0.021 | |
| KZS20464 | | Prostaglandin reductase | | PROSTAGLANDIN REDUCTASE 1 | -1.251 | 0.002 | |
| KZS21947 | | Arf-GAP with GTPase, ANK repeat and PH domain-containing protein | | CENTAURIN-GAMMA-1A | -1.234 | 0.043 | |
| KZS11463 | | Aquaporin-3 | | AQUAPORIN OR AQUAGLYCEROPORIN RELATED | -1.22 | 0.047 | |
| KZS04930 | | Zinc carboxypeptidase A 1 | | PEPTIDASE M14 CARBOXYPEPTIDASE A DOMAIN-CONTAINING PROTEIN | -1.203 | 0.025 | |
| KZS19264 | | UDP-glucuronosyltransferase 1-4 | | EG:EG0003.4 PROTEIN-RELATED | -1.167 | 0.028 | |
| KZS09322 | | Tyrosine-protein phosphatase non-receptor type | | MALTASE A1 | -1.159 | 0.009 | |
| KZS14409 | | Serine Protease | | AT07769P-RELATED | -1.12 | <0.001 | |
| KZS18300 | | Prostaglandin G/H synthase | | EGF-LIKE DOMAIN-CONTAINING PROTEIN | -1.076 | 0.093 | |
| KZS17514 | | Eukaryotic translation initiation factor 2-alpha kinase 1 | | INTERFERON-INDUCED, DOUBLE-STRANDED RNA-ACTIVATED PROTEIN KINASE | -1.069 | 0.01 | |
| KZR99511 | | Histone H1 | | HISTONE H1.0 | -1.057 | 0.057 | |
| KZS16131 | | serine threonine-protein kinase | | IRE1-RELATED | -1.054 | 0.079 | |
| KZS21268 | | Chitin deacetylase 9 | | FAMILY NOT NAMED | -1.03 | <0.001 | |
| Up in BHB |  | |  | |  | |  |
| KZS02589 | Uncharacterized protein | | 60S RIBOSOMAL PROTEIN L31 | | 15.633 | | 0.007 |
| KZS14798 | Zinc metalloproteinase nas-13 | | DISCOIDIN, CUB, EGF, LAMININ , AND ZINC METALLOPROTEASE DOMAIN CONTAINING | | 3.363 | | <0.001 |
| KZS21309 | Defective proboscis extension response | | DEFECTIVE PROBOSCIS EXTENSION RESPONSE 11, ISOFORM B-RELATED | | 1.143 | | 0.064 |
| Down in BHB |  | |  | |  | |  |
| KZS10907 | RVT_1, Reverse transcriptase (RNA-dependent DNA polymerase) | | REVERSE TRANSCRIPTASE | | -26.459 | | 0.002 |
| KZS10729 | Uncharacterized protein | |  | | -24.556 | | 0.005 |
| KZS20869 | ANK, Ankyrin repeats | | TRANSPOSASE | | -22.403 | | 0.02 |
| KZS19730 | Transcription factor Sox-7 | |  | | -21.639 | | 0.031 |
| KZS00976 | Uncharacterized protein | |  | | -21.53 | | 0.031 |
| snap_masked-falcon_000001F-processed-gene-41.107 | Chitin deacetylase 9 | |  | | -20.315 | | 0.064 |
| KZS03747 | Uncharacterized protein | | BEN DOMAIN-CONTAINING PROTEIN 6 | | -20.12 | | 0.071 |
| KZS05290 | cuticle protein | | CUTICULAR PROTEIN 50CB-RELATED | | -19.823 | | 0.071 |
| KZS03586 | Uncharacterized protein | |  | | -5.224 | | 0.002 |
| KZS15417 | Uncharacterized protein | | PROTEIN, PUTATIVE-RELATED | | -4.132 | | 0.012 |
| KZS06091 | Transport protein Sec23A | | PROTEIN TRANSPORT PROTEIN SEC23B | | -2.339 | | 0.064 |

| Up in NMN |  |  |  |  |
| --- | --- | --- | --- | --- |
| KZS03333 | ATP-dependent DNA helicase PIF1,EC=3.6.1.-/sw | ATP-DEPENDENT DNA HELICASE PIF1 | 5.48 | 0.023 |
| KZS21334 |  |  | 4.007 | 0.045 |
| KZS13203 | tartan | GLYCOPROTEIN LIG-1, PUTATIVE-RELATED | 3.628 | 0.01 |
| Down in NMN |  |  |  |  |
| KZS16723 | Complement C1q tumor necrosis factor-related protein 4 | C1Q DOMAIN-CONTAINING PROTEIN | -29.526 | <0.001 |

Table S5. Top GOs showing up- and down-regulation in response to caloric restriction. GO terms filtered to the most specific term.

| Up in CR |  |  |  |  |  |
| --- | --- | --- | --- | --- | --- |
| GO or pathway | ID | FET P-value | FDR | N genes | Expected |
| SCF-dependent proteasomal ubiquitin-dependent protein catabolic process | GO:0031146 | 0.0012 | 0.57 | 2 | 0.059 |
| oxidoreductase | PC00176 | 0.0029 | 0.33 | 3 | 0.293 |
| phospholipase | PC00186 | 0.0031 | 0.20 | 2 | 0.089 |
| carboxylic acid transmembrane transporter activity | GO:0046943 | 0.0042 | 0.22 | 2 | 0.102 |
| organonitrogen compound catabolic process | GO:1901565 | 0.0043 | 0.17 | 4 | 0.673 |
| calcium ion binding | GO:0005509 | 0.0071 | 0.23 | 2 | 0.132 |
| regulation of MAPK cascade | GO:0043408 | 0.0071 | 0.21 | 2 | 0.132 |
| Down un CR |  |  |  |  |  |
| extracellular matrix structural protein | PC00103 | 0.0019 | >1 | 5 | 0.922 |
| hydrolase | PC00121 | 0.0081 | >1 | 4 | 0.819 |
| metalloprotease | PC00153 | 0.02 | >1 | 4 | 1.058 |
| lipid transport | GO:0006869 | 0.022 | >1 | 2 | 0.239 |
| vesicle tethering complex | GO:0099023 | 0.028 | >1 | 2 | 0.273 |
